# Early cellular mechanism of type I interferon-driven susceptibility to tuberculosis

**DOI:** 10.1101/2022.10.06.511233

**Authors:** Dmitri I. Kotov, Ophelia V. Lee, Stefan A. Fattinger, Charlotte Langner, Jaresley V. Guillen, Joshua M. Peters, Andres Moon, Eileen M. Burd, Kristen C. Witt, Daniel B. Stetson, David L. Jaye, Bryan D. Bryson, Russell E. Vance

## Abstract

*Mycobacterium tuberculosis* (*Mtb*) causes 1.6 million deaths annually. Active tuberculosis correlates with a neutrophil-driven type I interferon (IFN) signature, but the cellular mechanisms underlying tuberculosis pathogenesis remain poorly understood. We found interstitial macrophages (IMs) and plasmacytoid dendritic cells (pDCs) are dominant producers of type I IFN during *Mtb* infection in mice and non-human primates, and pDCs localize near human *Mtb* granulomas. Depletion of pDCs reduces *Mtb* burdens, implicating pDCs in tuberculosis pathogenesis. During IFN-driven disease, we observe abundant DNA-containing neutrophil extracellular traps (NETs) known to activate pDCs. Cell type-specific disruption of the type I IFN receptor suggests IFNs act on IMs to inhibit *Mtb* control. Single cell RNA-seq indicates type I IFN-responsive cells are defective in their response to IFN*γ*, a cytokine critical for *Mtb* control. We propose pDC-derived type I IFNs act on IMs to drive bacterial replication, further neutrophil recruitment, and active tuberculosis disease.

## Introduction

*Mycobacterium tuberculosis* (*Mtb*), the causative agent of tuberculosis disease, caused 1.6 million deaths in 2021^1^. Treatment requires a minimum 4-6 month course of antibiotics, or up to 2 years for increasingly prevalent multi-drug resistant strains. Moreover, the only approved vaccine for *Mtb* has variable or no efficacy in adults^2^. The pathophysiology of tuberculosis remains poorly understood. The mouse model has been used to discover most of the host factors known to control tuberculosis in humans, including tumor necrosis (TNF) factor and interferon-*γ*^3–5^. Nevertheless, the use of mice has been criticized for poorly recapitulating key aspects of human disease^6^.

In humans, active tuberculosis disease is reproducibly associated with the induction of type I interferons (IFNs)^7–10^, a family of cytokines, including IFNα and IFNβ, that signal via the type I IFN receptor (IFNAR). Type I IFNs elicit a primarily anti-viral response that overlaps but is insufficient to recapitulate the protective anti-*Mtb* response elicited by IFN*γ*. Infection with type I IFN-inducing viruses is associated with worse *Mtb* infection outcomes in humans. For example, influenza infection correlates with an increased risk of death among patients with pulmonary tuberculosis, and infants with cytomegalovirus have an increased risk of tuberculosis disease^11–13^. These human studies do not prove that the correlation between virus infection and *Mtb* disease is due to the induction of type I IFNs. However, a causal role for type I IFNs in driving human tuberculosis is supported by the finding that a partial loss-of-function mutation in *IFNAR* is associated with *Mtb* resistance in humans^14^. Moreover, a type I IFN response has the potential to be a driver of progression to active *Mtb* in humans, as type I IFNs can antagonize the critical, protective IFN*γ* response during mycobacterial infection^5,15,16^.

Mice are able to model the virus-induced susceptibility to *Mtb* seen in humans, with chronic lymphocytic choriomeningitis virus (LCMV), acute LCMV, pneumonia virus of mice, and influenza A virus co-infection with *Mtb* all exacerbating tuberculosis disease^17–21^. Type I IFNs were shown to drive the loss of *Mtb* control during the acute LCMV and influenza co-infections^17,18^. However, co-infection studies make it challenging to determine the cellular mechanism behind *Mtb* susceptibility, as perturbations such as type I IFN receptor blockade simultaneously impact viral and bacterial control. Therefore, an ideal platform for studying how type I IFN induces loss of *Mtb* control would be mice in which *Mtb* infection is itself sufficient to elicit the exacerbated type I IFN response that is seen in humans with active TB. However, C57BL/6 (B6) mice, the most used model for *Mtb* infection, generate a weak type I IFN response in response to *Mtb* infection; indeed, *Ifnar1* deletion from unmanipulated B6 mice does not consistently impact bacterial *Mtb* burdens in the lungs^22–24^. Investigators have circumvented this issue by intranasal injection of B6 mice with polyI:C, an IFN-inducing viral mimic^9,25,26^. Such studies demonstrate a convincing causal link between type I IFNs and *Mtb* susceptibility, and have shown that a major detrimental effect of type I IFNs is to impair interleukin-1-dependent control of *Mtb*^26^. However, despite these advances, it has been challenging to decipher the cellular mechanisms underlying the detrimental effects of type I IFN on bacterial control, and in particular, the cells required to produce or respond to type I IFNs to mediate *Mtb*-susceptibility remain unknown.

Unlike B6 mice, C3H and 129 mice exhibit a type I interferon-driven susceptibility to *Mtb*^27^, but there are limited genetic tools in these mouse strains, making mechanistic studies difficult. We recently discovered that congenic B6 mice with the ‘super susceptibility to tuberculosis 1’ region from C3H mice (i.e., B6.*Sst1^S^*mice)^28,29^ are highly susceptible to *Mtb* infection due to their strong type I IFN response. *Ifnar1* deletion fully rescues the susceptibility of B6.*Sst1^S^* mice at early timepoints and enhances survival^30^. We recently identified *Sp140* as the gene within the *Sst1* genetic interval that controls *Mtb* susceptibility, and confirmed that the early *Mtb* susceptibility of *Sp140*^−/−^ mice is also rescued by *Ifnar1* deletion^31^. As our *Sp140*^−/−^ mice were generated on a pure C57BL/6J background, this mouse model enables mechanistic genetic studies of the type I IFN response during *Mtb* infection.

In the present study, we leveraged *Sp140*^−/−^ mice to determine the cellular mechanisms by which type I IFN drives *Mtb* susceptibility. Single cell RNA-sequencing (scRNA-seq) identified interstitial macrophages (IMs) as a major type I IFN producer. A sensitive genetic reporter of type I IFN production corroborated the scRNA-seq findings, and also revealed that plasmacytoid dendritic cells (pDCs) are an additional source of type I IFN during *Mtb* infection. Type I IFN production by pDCs appears to drive disease since pDC depletion rescued the susceptibility of *Sp140*^−/−^ mice. Cell-type specific deletion of *Ifnar1* demonstrated that type I IFN confers susceptibility by acting on IMs. We developed transcriptional signatures that distinguish the response elicited by type I IFN from that elicited by IFNγ. Application of these signatures to our scRNA-seq data indicates that *Mtb*-infected type I IFN-responsive IMs in the lungs are impaired in their responses to IFN*γ*^16,32,33^. Loss of bacterial control in *Sp140*^−/−^ mice leads to an influx of neutrophils and abundant production of DNA-rich neutrophil extracellular traps (NETs), ligands known to promote type I IFN production by pDCs^34,35^. Our findings suggest a new model of tuberculosis pathogenesis in which type I IFNs drive an initial loss of bacterial control, possibly by impairing responses to IFN*γ*, which in turn initiates a positive feedback loop of NET production and type I IFN expression by pDCs, leading to uncontrolled bacterial replication and active tuberculosis disease.

## Results

### Myeloid cells harbor Mtb in Sp140^−/−^ mice

As we have previously demonstrated, the susceptibility of *Sp140*^−/−^ mice to *Mtb* infection is driven by type I IFN, and mirrors the correlation between type I IFN and active tuberculosis disease seen in humans. Genetic or antibody-mediated depletion of IFNAR fully rescues the enhanced susceptibility of *Sp140*^−/−^ animals (**Fig. 1A**)^31^. To better characterize the immune response of *Sp140*^−/−^ mice to *Mtb*, we infected mice with *Mtb* harboring a Wasabi fluorescent protein expression plasmid^36^, which permits robust detection of *Mtb*-infected cells by flow cytometry. With this approach, we were able to clearly identify *Mtb*-harboring cells in mouse lungs 25 days post-infection (**Fig. 1B**). The vast majority of the bacteria isolated from these lungs were highly fluorescent, indicating that the plasmid is stably maintained for at least 25 days. The number of *Mtb*-infected cells detected by flow cytometry strongly correlates with lung *Mtb* CFU (R^2^ = 0.63), indicating that flow cytometry reliably reports overall lung bacterial burden 25 days after *Mtb* infection (**Fig. 1C**). We classified lung immune cells as CD11b^+^ Ly6G^+^ neutrophils, CD64^+^ MerTK^+^ Siglec F^+^ alveolar macrophages (AMs), CD64^+^ MerTK^+^ Siglec F^−^ CD11b^+^ IMs, CD64^+^ MerTK^−^ CD11b^+^ monocytes, CD64^−^ CD11c^+^ MHCII^hi^ CD11b^−^ conventional type I dendritic cells (cDC1), or CD64^−^ CD11c^+^ MHCII^hi^ CD11b^+^ conventional type II dendritic cells (cDC2) and assessed the abundance of these cells at day 25 after *Mtb* infection in *Sp140*^−/−^ and *Sp140*^−/−^ *Ifnar1*^−/−^ mice (**Supplementary Fig. 1**). The lungs of infected *Sp140*^−/−^ animals contained significantly more neutrophils, IMs, and monocytes as compared to *Sp140*^−/−^ *Ifnar1*^−/−^ mice, with no difference in the number of AMs and a reduction in the number of cDC2 (**Fig. 1D**). Over 90% of the infected cells were myeloid cells (**Fig. 1B, 1E**), in line with previous reports^37,38^. Consistent with their higher abundance in infected *Sp140*^−/−^ lungs, neutrophils comprised a considerably larger percentage and absolute number of the *Mtb*-infected cells in *Sp140*^−/−^ mice compared to *Sp140*^−/−^ *Ifnar1*^−/−^ animals (**Fig. 1E, 1F**). There were also more infected IMs and monocytes in *Sp140*^−/−^ mice compared to *Sp140*^−/−^ *Ifnar1*^−/−^ animals, in line with the overall increase in these immune populations in the lungs of infected *Sp140*^−/−^ mice (**Fig. 1D, 1F**). Importantly, we did not observe alterations in the immune compartment of uninfected *Sp140*^−/−^ mice, implying grossly normal hematopoietic development in these mice (**Supplemental Fig. 2**)^39–41^. However, the exact mechanisms causing the infection-induced differences in myeloid cells from *Sp140*^−/−^ mice was unclear, and thus required more in-depth profiling of the myeloid compartment.

**Figure 1.**
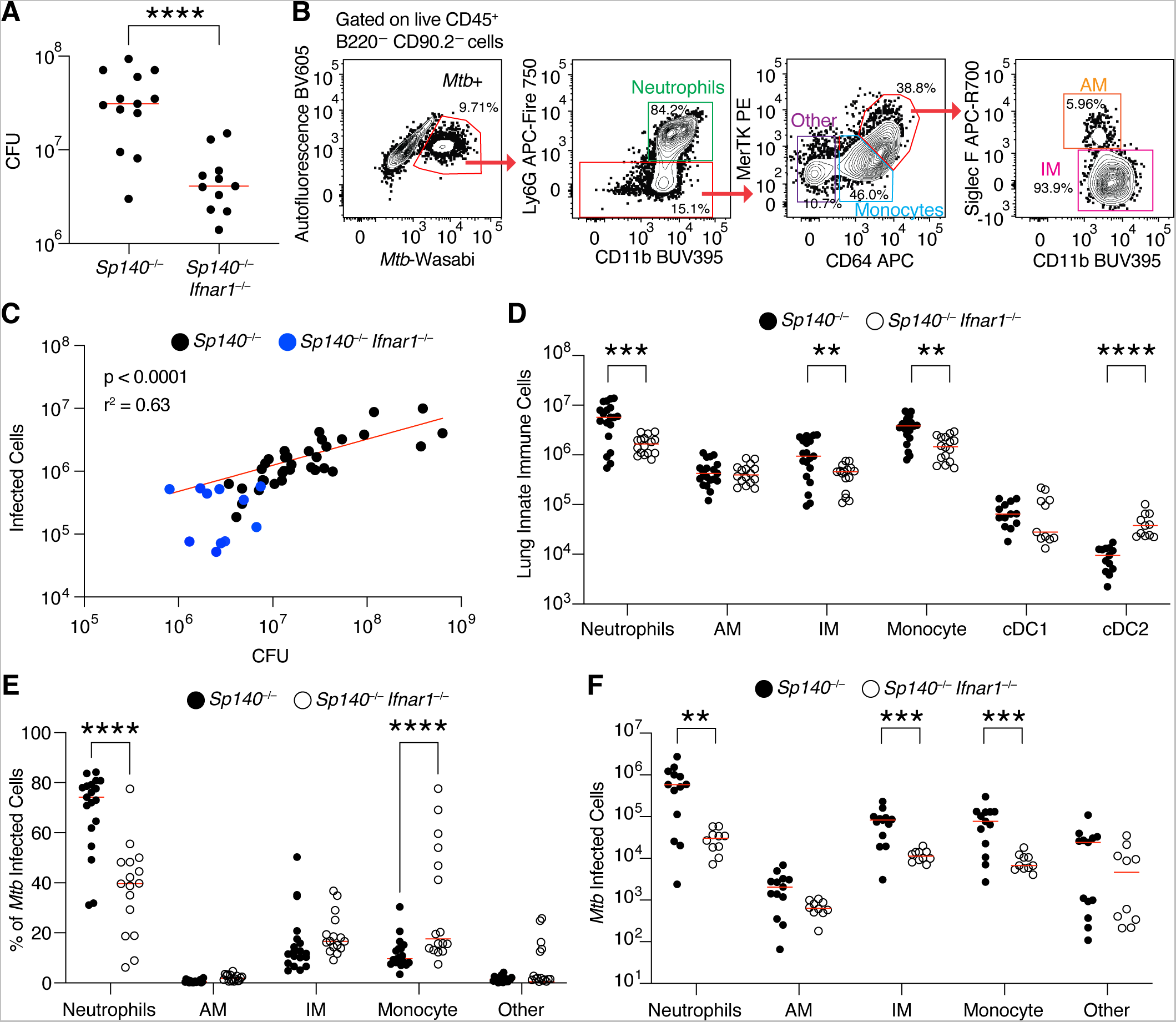
Myeloid cells are the dominant *Mtb* harboring cells in *Sp140*^−/−^ and *Sp140*^−/−^ *Ifnar1*^−/−^ mice. (**A**) Colony forming units (CFU) of *Mtb* in the lungs of *Sp140*^−/−^ (n = 13) and *Sp140*^−/−^ *Ifnar1*^−/−^ (n = 11) mice 25 days post-infection. (**B**) Representative flow cytometry plots of an *Mtb*-infected *Sp140*^−/−^ mouse lung gated on live CD45^+^ B220^−^ CD90.2^−^ cells to identify *Mtb*-infected cells, subset into neutrophils (green; Ly6G^+^ CD11b^+^), other cells (purple; Ly6G^−^ CD64^−^ MerTK^−^), monocytes (blue; Ly6G^−^ CD64^+^ MerTK^low^), alveolar macrophages (AMs; orange; Ly6G^−^ CD64^+^ MerTK^high^ Siglec F^+^), and interstitial macrophages (IMs; pink; Ly6G^−^ CD64^+^ MerTK^high^ Siglec F^−^). (**C**) Correlation between the number of infected cells identified by flow cytometry of total lung digests gated as in panel B to the colony forming units from the same infected lung (black; *Sp140*^−/−^ and blue; *Sp140*^−/−^ *Ifnar1*^−/−^ combined; n = 45). The red line indicates a nonlinear regression of the displayed data. (**D**) Number of innate immune cells by cell type in the lungs of *Mtb*-infected *Sp140*^−/−^ (n = 19; closed circles) and *Sp140*^−/−^ *Ifnar1*^−/−^ mice (n = 16; open circles). (**E**) Frequency and (**F**) number of immune cell populations of *Mtb*-infected cells in *Sp140*^−/−^ (n = 12-19; closed circles) and *Sp140*^−/−^ *Ifnar1*^−/−^ mice (n = 10-15; open circles). Lungs were analyzed for the depicted experiments 24-26 days after *Mtb* infection. The bars in (A), (D), (E), and (F) represent the median. Pooled data from two or three independent experiments are shown. A linear regression performed on log transformed data was used to calculate significance and R^2^ for (C). An unpaired t test was used to determine significance for (A), a two-way ANOVA with Sidak’s multiple comparisons test was used to calculate significance for (D), (E), and (F). **p < 0.01, ***p < 0.001, ****p < 0.0001.

### Macrophages and neutrophils exhibit a variety of activation states during Mtb infection

To further characterize the *Mtb*-infected myeloid cells and dissect the cellular mechanism of type I IFN driven *Mtb* susceptibility, we compared B6 and *Sp140*^−/−^ innate immune responses by performing scRNA-seq on myeloid cells from *Mtb*-infected or uninfected lungs 25 days after infection. For this experiment, CD64^+^ and Ly6G^+^ cells were magnetically enriched, sort purified, and processed for library generation with the 10X Genomics platform (**Fig. 2A**). To ensure proper cell clustering, mRNA transcripts and protein expression for select lineage markers were simultaneously measured by CITE-seq (**Supplementary Fig. 3**)^42^. Resulting datasets were analyzed with Seurat V4, and Weighted Nearest Neighbor (WNN) analysis was used to cluster cells based on mRNA and protein expression^43^. Analyzed datasets were then visualized by uniform manifold approximation and projection (UMAP) reduction on the WNN clustered data (wnnUMAP)^44^. The resulting dataset consists of 6,604 B6 and 13,668 *Sp140*^−/−^ cells, almost exclusively consisting of myeloid cells (**Fig. 2B**). Major cell types were annotated based on protein or mRNA expression of lineage defining markers, such as Siglec F protein expression for identifying AMs (**Supplementary Fig. 4**)^45^. Individual clusters within a major cell type, like the 10 clusters of neutrophils, were annotated based on expression of maturation and activation markers (**Supplementary Fig. 5**)^46^. Each cluster is represented in the B6 and *Sp140*^−/−^ datasets, however the proportions of some clusters are altered between genotypes. Most notably, the ratio of IFN stimulated gene (ISG)^+^ IM to ISG^−^ IM was higher in the *Sp140*^−/−^ mice, as compared to B6, as expected from the exacerbated type I IFN response in *Sp140*^−/−^ mice (**Fig. 2B**). The largest changes in composition are seen when comparing cells from naïve lungs to bystander and *Mtb*-infected cells from *Mtb*-infected lungs (**Fig. 2B**). For example, AMs are relatively abundant in naïve lungs but are rare among the *Mtb*-infected cells 25 days post-infection, as also seen by flow cytometry (**Fig. 1F, 2B**). To establish whether the differences in immune response between *Sp140*^−/−^ and B6 mice occurred in response to *Mtb* infection, we compared naïve *Sp140*^−/−^ and B6 lungs by scRNA-seq. Consistent with the normal cellular profile of naïve *Sp140*^−/−^ mice (**Supplementary Fig. 2**), fewer than 10 differentially expressed genes were identified between the two genotypes in AMs, IMs, monocytes, and neutrophils (**Supplementary Fig. 6**). These results suggest that the immune compartment of B6 and *Sp140*^−/−^ mice is highly similar at baseline, and the type I IFN-driven changes in the genotypes occur after *Mtb* infection.

**Figure 2.**
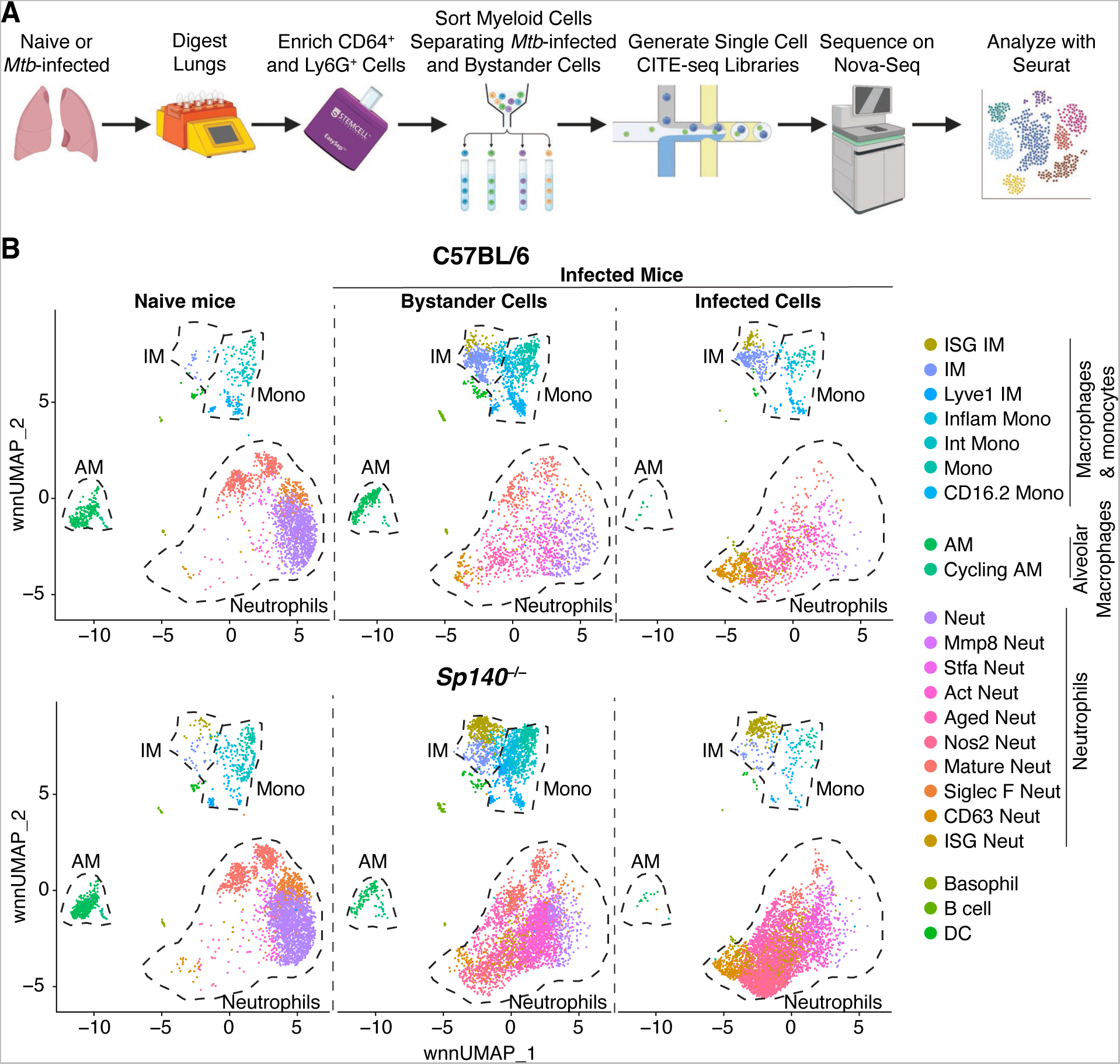
Single cell RNA-sequencing analysis of B6 and *Sp140*^−/−^ myeloid cells from *Mtb*-infected and naïve lungs. CITE-seq was used to perform scRNA-sequencing analysis to integrate transcriptomic and protein expression of single cells, as described further in Supplementary Fig. 3. (**A**) Model of the processing steps involved in generating the scRNA-sequencing dataset. (**B**) wnnUMAP plot depicting unbiased clustering of myeloid cells in B6 and *Sp140*^−/−^ *Mtb*-infected and naïve lungs (n = 10 lungs; n= 20,272 cells). The wnnUMAP plot depicts cells from naïve mice, the bystander cells from *Mtb*-infected mice, and the *Mtb*-infected cells from *Mtb*-infected mice. Lungs were analyzed for the depicted experiment 25 days after *Mtb*-Wasabi infection. For each genotype, n = 3 for the infected lung samples and n = 2 for the naïve lung samples.

### Bystander pDCs and IMs are the primary sources of type I IFN during Mtb infection

To determine the cellular mechanism of type I IFN-driven *Mtb* susceptibility, we first sought to identify which cells produce type I IFN following infection. In general, our scRNAseq analysis revealed that very few cells were *Ifnb1* positive, which may reflect a lack of sensitivity of scRNAseq, and/or the transient and stochastic expression pattern of this gene (**Fig. 3A**)^47–50^. *Mtb* infection resulted in increased expression of *Ifnb1* in infected and bystander mononuclear phagocytes, with a slight bias towards *Ifnb1* production by IMs compared to monocytes, and no production by AMs (**Fig. 3A**). While there was no major difference in the cell types producing *Ifnb1* between B6 and *Sp140*^−/−^ cells, a greater number and frequency of *Sp140*^−/−^ cells expressed *Ifnb1* compared to B6 cells (**Fig. 3B**). Additionally, *Ifnb1* expressing cells in *Sp140*^−/−^ mice trended towards a higher per cell expression of *Ifnb1* than B6 cells, but this difference was not significant (**Fig. 3C**). Consistent with these findings, a prior scRNA-seq study of *Mtb*-infected and naïve lungs from non-human primates largely mirrors our findings in mice^51^. Our analyses of these data indicate that IMs were also the dominant *IFNB1*-expressing cells in non-human primates with active tuberculosis, and IMs did not express *IFNB1* in naïve or latently infected lungs (**Fig. 3D**). These results suggest that mice faithfully recapitulate the *Mtb*-induced type I IFN production seen in non-human primates.

**Figure 3.**
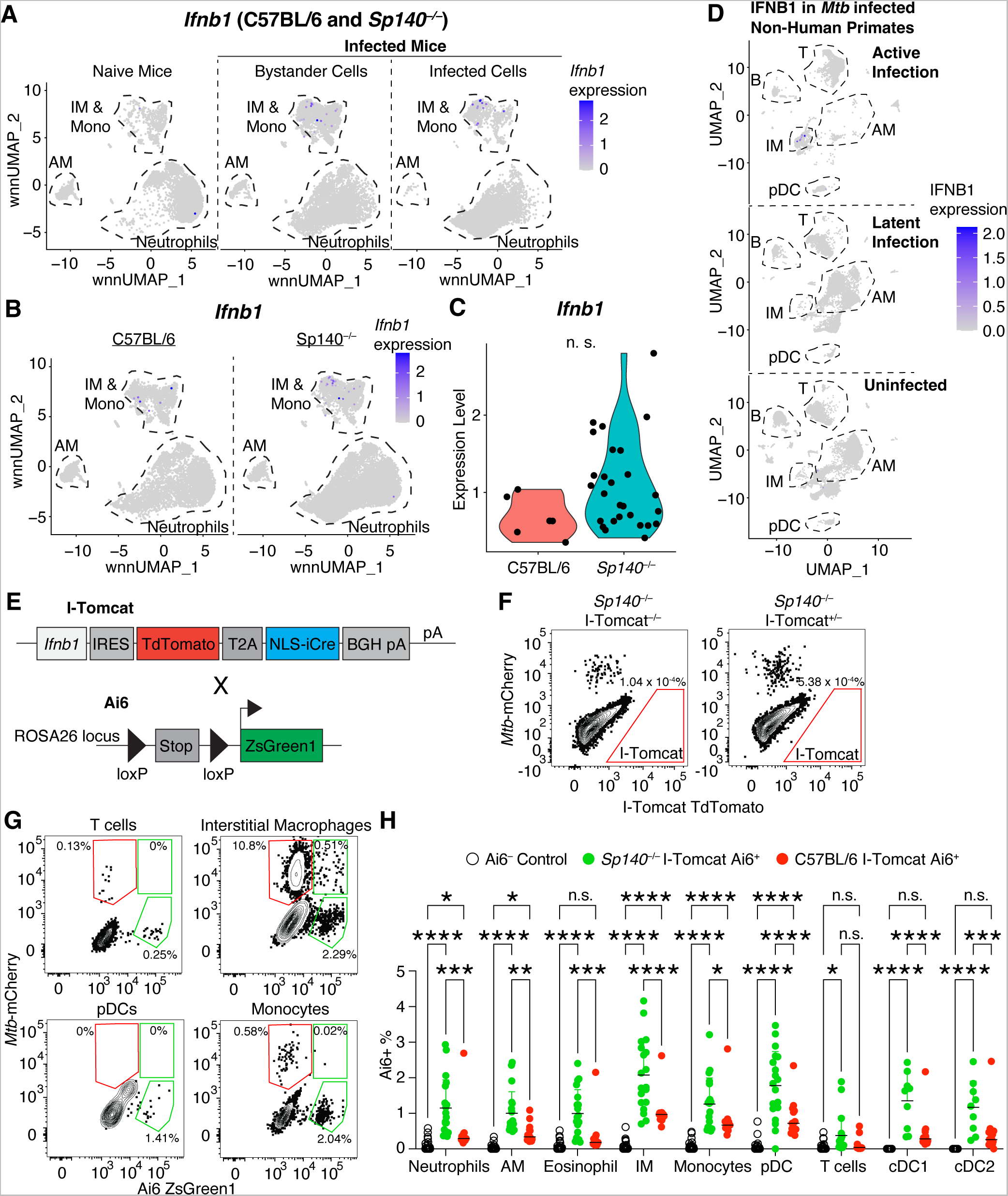
Bystander pDCs, IMs, and monocytes are the primary IFN-β producers in mice and non-human primates. (**A**) wnnUMAP plot of *Ifnb1* expression in myeloid cells from naïve mice, bystander myeloid cells from infected mice, and *Mtb*-infected myeloid cells from infected mice (B6 and *Sp140*^−/−^ combined). (**B**) wnnUMAP plot of *Ifnb1* expression in myeloid cells (combined infected and bystander) from B6 and *Sp140*^−/−^ mice. (**C**) Violin plot depicting level of *Ifnb1* expression in B6 and *Sp140*^−/−^ cells that express *Ifnb1*. (**D**) Analysis of GSE149758 scRNA-seq data from Esaulova E., *et al*. 2021. UMAP depicting *IFNB1* expression in cells from non-human primates with active *Mtb* infection, latent *Mtb* infection, or that are uninfected. (**E**) Schematic representation of the genetic structure of I-Tomcat mice and Ai6 mice. Cre expression permanently deletes the loxP-flanked STOP cassette resulting in constitutive ZsGreen expression. (**F**) Representative flow cytometry plot indicating lack of detectable TdTomato expression in immune cells. (**G**) Representative flow cytometry plots of ZsGreen expression and *Mtb*-mCherry detection in T cells, IMs, pDCs, and monocytes. (**H**) Frequency of Ai6 expressing cells in lung immune cells from Ai6^−^ control (n = 34; open circles), *Sp140*^−/−^ I-Tomcat Ai6 (n = 19; green circles), and I-Tomcat Ai6 (n = 15; red circles) mice. The bars in (H) represent the median. Pooled data from four independent experiments are shown in (H). Lungs were analyzed for the depicted experiments 25 days after *Mtb* infection. Statistical significance in (C) was calculated by non-parametric Wilcoxon rank sum test with Bonferroni correction and in (H) by one-way ANOVA with Tukey’s multiple comparison test. *p < 0.05, **p < 0.01, ***p < 0.001, ****p < 0.0001.

The type I IFN producers identified in the scRNA-seq datasets were validated using a genetic reporter of type I IFN production, called I-Tomcat mice (from Dan Stetson, manuscript in preparation). These mice express TdTomato and Cre downstream of *Ifnb1*; therefore, any cell that expresses *Ifnb1* will also express TdTomato and Cre (**Fig. 3E**). While TdTomato expression was sufficient to identify *Ifnb1* expression by I-Tomcat bone marrow-derived macrophages following *in vitro* stimulation with poly I:C (**Supplementary Fig. 7A**), TdTomato^+^ cells were not detected 25 days after *Mtb* infection (**Fig. 3F**). The ability to detect TdTomato^+^ cells was not improved by using I-Tomcat homozygous mice, examining an earlier timepoint of 19 days post-infection, or by gating on specific immune populations such as IMs (**Supplementary Fig. 7B, 7C, 7D**). Even though type I IFN drives the *Mtb* susceptibility of *Sp140*^−/−^ mice 25 days post-infection, it is unclear when the type I IFN production occurs (**Fig. 1A**). It is possible that type I IFN is an early and/or transient event, which would be missed by analyzing a single timepoint with the I-Tomcat mice. To address this issue, we crossed I-Tomcat mice with the Ai6 Cre reporter mouse line to generate I-Tomcat Ai6 mice on both the B6 and *Sp140*^−/−^ backgrounds (**Fig. 3E**)^52^. In these mice, any cell that has ever expressed *Ifnb1* will constitutively express ZsGreen, allowing for their sensitive detection; importantly, however, the levels of ZsGreen do not report the levels of *Ifnb1* expression. *Mtb*-infected I-Tomcat Ai6 mice clearly contained populations of reporter-positive myeloid cells, while demonstrating low background among cell populations that are not expected to be reporter positive (e.g., ~0.1% of T cells were Ai6^+^) (**Fig. 3G**). Consistent with the scRNAseq analysis, IMs and monocytes were the primary *Ifnb1* expressing cells in B6 and *Sp140*^−/−^ mice (**Fig. 3H**, **Supplementary Fig. 8**). Interestingly, *Sp140*-deficient mice exhibited elevated Ai6^+^ expression frequency in all cell types, suggesting SP140 broadly modulates the sensitivity for inducing *Ifnb1* expression (**Fig. 3H**). In addition to corroborating the scRNA-seq data, the I-Tomcat mice also identified pDCs as a major type I IFN producing cell population. Lung pDCs are very rare and were therefore not represented in our scRNA-seq dataset, demonstrating the power of using genetic reporters to study rare events and cell populations. Despite their scarcity, pDCs are known to be extremely robust producers of type I IFNs on a per cell basis^53^.

While we expected IMs to be a major type I IFN producing population given the scRNA-seq results, we were surprised that the majority of the Ai6^+^ IMs were *Mtb*^−^ and most *Mtb*^+^ IMs were Ai6^−^ (**Fig. 3G**). These results suggest that direct infection of IMs is neither required nor sufficient for IFN-β production. To examine this phenomenon in greater detail, we performed confocal microscopy and histo-cytometry analysis of *Mtb*-infected I-Tomcat Ai6 and *Sp140*^−/−^ I-Tomcat Ai6 lungs^54,55^. While lesions of diseased tissue were clearly identifiable in I-Tomcat Ai6 mice, the size and myeloid cell influx into the diseased tissue were greatly exacerbated in *Sp140*^−/−^ I-Tomcat Ai6 (**Fig. 4A**). Additionally, Ai6 expressing cells were identifiable throughout the lungs, with an increased propensity to localize in diseased rather than healthy tissue (**Fig. 4A**, **Supplementary Fig. 9**). Within diseased tissue, Ai6 expressing cells were primarily located near *Mtb* harboring cells in I-Tomcat Ai6 and *Sp140*^−/−^ I-Tomcat Ai6 lungs (**Fig. 4B**). Similar to the flow cytometry results, SIRPɑ^+^ cells, which are primarily macrophages in *Mtb*-infected lungs as they are ~100 fold more abundant than SIRPɑ expressing cDC2s, were a major Ai6 expressing cell population (Fig. 1D, 4B, 4C). The SIRPɑ^+^ macrophages expressed Ai6 at a higher frequency than CD4^+^ T cells in the diseased tissue but not healthy tissue (**Fig. 3H, 4B, 4C**). Direct infection by *Mtb* was not a major driver of IFN-β expression, as ~2-3% of infected macrophages were Ai6^+^ and ~12-15% of Ai6^+^ cells were *Mtb*-infected, in line with the frequencies seen in IMs by flow cytometry (**Fig. 3H, 4D**). These results suggest that IM localization to *Mtb* rich regions provides the activating signals required for IFN-β expression, while direct infection of IMs is not required for IFN-β expression.

**Figure 4.**
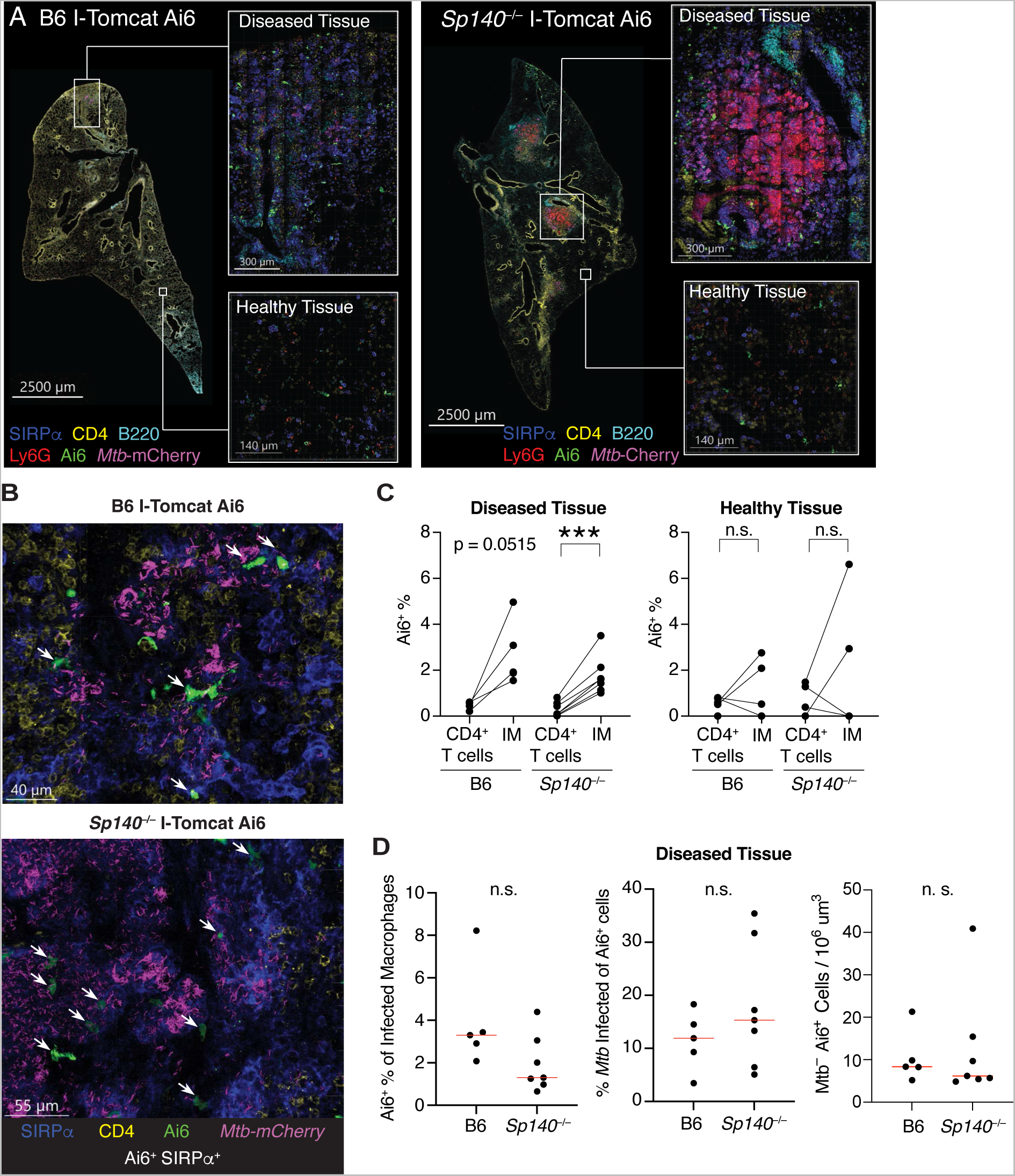
Cells producing IFN-β are enriched in diseased tissue, but only a minority harbor *Mtb*. (**A**) Representative images of *Mtb*-infected I-Tomcat Ai6 and *Sp140*^−/−^ I-Tomcat Ai6 lung sections stained for SIRPɑ (dark blue), CD4 (yellow), B220 (teal), Ly6G (red), Ai6 (green), and *Mtb*-expressed mCherry (magenta). Inset images depict higher magnification of diseased and healthy tissue for both genotypes. (**B**) Representative images of Ai6^+^ cell localization near *Mtb* in the diseased portions of I-Tomcat Ai6 and *Sp140*^−/−^ I-Tomcat Ai6 lungs. Sections were stained with SIRPɑ (dark blue), CD4 (yellow), Ai6 (green), and *Mtb*-expressed mCherry (magenta). White arrows indicate cells co-expressing Ai6 and SIRPɑ. (**C**) Image quantification of the frequency of Ai6 expression in CD4^+^ T cells and SIRPɑ^+^ IMs in the diseased and healthy tissue of B6 I-Tomcat Ai6 (n = 5) and *Sp140*^−/−^ I-Tomcat Ai6 (n = 7) lungs. (**D**) Image quantification of the frequency of Ai6 expression among *Mtb*-infected macrophages, frequency of *Mtb* infection among Ai6^+^ cells, and number of uninfected Ai6^+^ cells for B6 I-Tomcat Ai6 (n = 5) and *Sp140*^−/−^ I-Tomcat Ai6 (n = 7) lungs. All samples were analyzed 25 days after *Mtb* infection. Pooled data from two independent experiments are shown in (C) and (D). Statistical significance in (C) and (D) was calculated with an unpaired t test. ***p < 0.001.

### pDCs significantly contribute to the Mtb susceptibility of Sp140^−/−^ animals

While pDCs have a well-established role in anti-viral immunity in the lung, limited work has assessed their contribution during *Mtb* infection^56,57^. pDCs may have been previously overlooked because of their scarcity in the lung. Indeed, we observe only ~20,000 pDCs in the lungs of naïve B6 and *Sp140*^−/−^ mice, but this number increases 10-fold following *Mtb* infection and is significantly higher in *Mtb*-infected *Sp140*^−/−^ than B6 mice (**Fig. 5A**). Despite their scarcity, pDCs can have major effects due to the extremely high levels of interferons produced per cell^53^. Consistent with our finding that pDCs are a major type I IFN producer in *Mtb*-infected mouse lungs, Khader and colleagues described the presence of pDCs in lungs of non-human primates with active pulmonary TB^51^. However, the lack of genetic tools in non-human primates precluded functional studies of pDCs during TB. Therefore, we decided to take advantage of our experimentally tractable mouse model to assess whether pDCs affect *Mtb* control. The contribution of pDCs was initially tested by depleting pDCs using an anti-BST2 antibody^58–60^. This strategy efficiently depleted pDCs and resulted in a partial rescue of *Mtb* control in *Sp140*^−/−^ mice (**Fig. 5B, 5C**). However, BST2 is known to be upregulated by cells other than pDCs in inflammatory environments, thus antibody depletion could have been protective against *Mtb* by depleting non-pDC cells^59^. We therefore also tested the contribution of pDCs by using a genetic pDC depletion strategy in which we crossed our *Sp140*^−/−^ mice with mice expressing the diphtheria toxin receptor (DTR) downstream of the human BDCA2 promoter (pDC-DTR)^61^. We used *Sp140*^+/−^ pDC-DTR littermates as wild-type controls since a single copy of *Sp140* is sufficient to restore wild-type control of *Mtb*. DT administration efficiently ablated pDCs in *Sp140*^−/−^ and *Sp140*^+/−^ mice, with the depletion specifically affecting pDCs (**Fig. 5D**, **Supplementary Fig. 10**). pDC-DTR depletion of pDCs was able to fully rescue bacterial control in *Sp140*^−/−^ mice, while depletion in *Sp140*-sufficient animals (that do not exhibit an exacerbated type I IFN response) did not affect lung bacterial burden, as expected (**Fig. 5E**). Additionally, pDC-DTR depletion of pDCs in *Sp140*^−/−^ mice reduced expression of type I IFN stimulated genes to the level of *Sp140*-sufficient animals, while rescuing expression of the type II IFN stimulated gene *H2-Ab1* (**Fig. 5F, 5G**). These results demonstrate a novel contribution by pDCs in limiting *Mtb* control in animals with a hyper type I IFN response.

**Figure 5.**
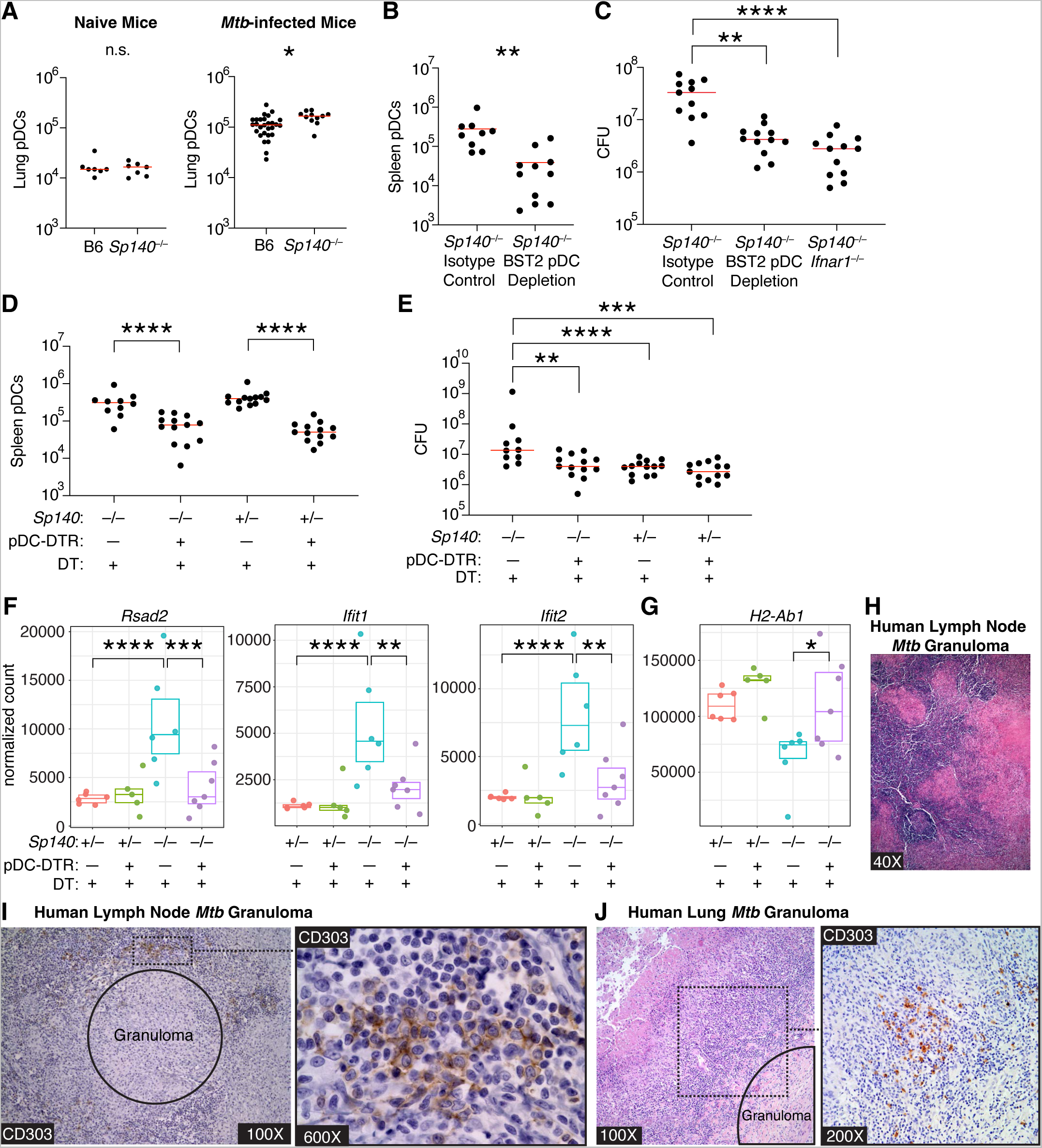
pDC depletion reduces *Mtb* burdens in *Sp140*^−/−^ mice, and pDCs are present in the lymphocytic cuff surrounding granulomas in *Mtb*-infected human lymph nodes and lungs. (**A**) Number of lung pDCs in B6 and *Sp140*^−/−^ mice in naïve (n = 7) and *Mtb*-infected mice (n = 11-28). (**B**) Number of splenic pDCs and (**C**) bacterial burden in *Sp140*^−/−^ mice that received isotype or pDC depleting anti-BST2 antibody from days 12 to 24 post-infection (n = 9-12). (**D**) Number of splenic pDCs, (**E**) bacterial burden, (**F**) lung expression of *Rsad2*, *Ifit1*, and *Ifit2* as representative type I IFN-stimulated genes, and (**G**) lung expression of *H2-Ab1* as a representative type II IFN-stimulated gene in *Sp140*^−/−^ pDC-DTR mice or *Sp140*^−/−^ mice controls that received DT from days 12 to 24 after infection (n = 10-13 for (D) and (E); n = 5-7 for (F) and (G)). (**H**) Representative hematoxylin and eosin or (**I**) anti-CD303 (brown) and hematoxylin staining on serial sections of *Mtb*-infected human lymph nodes (n = 8). (**J**) Representative hematoxylin and eosin and anti-CD303 (brown) and hematoxylin staining on serial sections of *Mtb*-infected human lung samples (n = 8). Mouse lungs were harvested 25 days post-infection. The bars in (A), (B), (C), (D), and (E) represent the median. Pooled data from two independent experiments are shown in (A), (B), (C), (D), and (E). Statistical significance was calculated by one-way ANOVA with Tukey’s multiple comparison test for (C), (D), and (E) and by an unpaired t test for (A) and (B). *p < 0.05, **p < 0.01, ***p < 0.001, ****p < 0.0001.

While our results demonstrate a role for pDCs during *Mtb* infection in mice, and the Khader lab identified a correlation between pDCs and active *Mtb* in non-human primates, the role of pDCs during human *Mtb* infection has yet to be examined^51^. Therefore, we analyzed human lung and lymph node biopsies taken from *Mtb* culture-positive patients for the presence of pDCs near *Mtb* granulomas (**Fig. 5H**). Based on CD303 and CD123 staining, pDCs localized to the lymphocytic cuff surrounding *Mtb* granulomas in human lungs and lymph nodes (**Fig. 5I, 5I**, **Supplementary Fig. 10**). Of the 8 patient samples analyzed, 5 lung samples and 7 lymph node samples had pDCs in the same 400× field as an *Mtb* granuloma (**Supplementary Table 1**). The majority of the pDCs in the lung samples were distributed as individual cells, while lymph node pDCs were primarily grouped together in clusters of over 20 cells or scattered individually (**Supplementary Table 1**). These results demonstrate that pDCs, though generally an extremely rare cell population, are nevertheless located near *Mtb*-infected cells in granulomas in human lung and lymph nodes. These results, along with our results in mice and previous studies in non-human primates^51^, implicate pDCs as a plausible source of type I IFN that drives active tuberculosis in humans.

To understand why pDCs contribute to the susceptibility of *Sp140*^−/−^ but not *Sp140*-sufficient animals, we examined the availability of ligands that might potentially activate pDCs to produce type I IFNs. We focused on DNA-rich neutrophil extracellular traps (NETs) as a potential pDC-activating ligand because extracellular DNA is known to activate pDCs via TLR9, and NETs have been described as a stimulus for type I interferon production by pDCs in mice and humans in the context of autoimmunity^34,35,62^. Additionally, another *Mtb* susceptible mouse model with a hyper type I interferon response identified the presence of NETs in the lungs of susceptible mice and humans with active *Mtb* disease^63^. We first assessed whether neutrophils were enriched in *Mtb*-infected lungs and found a 5-fold increase in neutrophils in *Sp140*^−/−^ relative to *Sp140*^−/−^ *Ifnar1*^−/−^ mice (**Fig. 6A**). Additionally, neutrophil depletion partially rescued the susceptibility of *Sp140*^−/−^ mice at lower and higher bacterial burdens (**Fig. 6B, 6C**, **Supplementary Fig. 11**). Given the importance of neutrophils, we assessed NET production in *Sp140*^−/−^ and B6 mice by staining for citrullinated H3 in the lungs of *Mtb*-infected mice (**Fig. 6D**). *Sp140*^−/−^ mice had over a 100-fold increase in NET staining as compared to B6 animals, indicating that the lungs of *Sp140*^−/−^ mice harbor substantially more ligand to activate type I interferon production by pDCs as compared to wild-type hosts (**Fig. 6E**). As NETs are a source of nucleic acids, we hypothesized that pDCs would sense the NETs through endosomal TLRs. In line with this prediction, deletion of *Unc93b1*, which is an essential chaperone required for TLR7 and TLR9 function, partially rescued the *Mtb* susceptibility of *Sp140*^−/−^ mice (**Fig. 6G**). By contrast, deletion of *Ticam1,* which encodes for TRIF, an adapter molecule critical for type I IFN production downstream of TLR3 and TLR4, had no effect on bacterial control in *Sp140*^−/−^ mice (**Fig. 6F**). Together, these data suggest a model in which extracellular nucleic acid, potentially from NETs, is sensed by endosomal TLRs triggering pDC production of type I IFNs (**Fig. 6H**).

**Figure 6.**
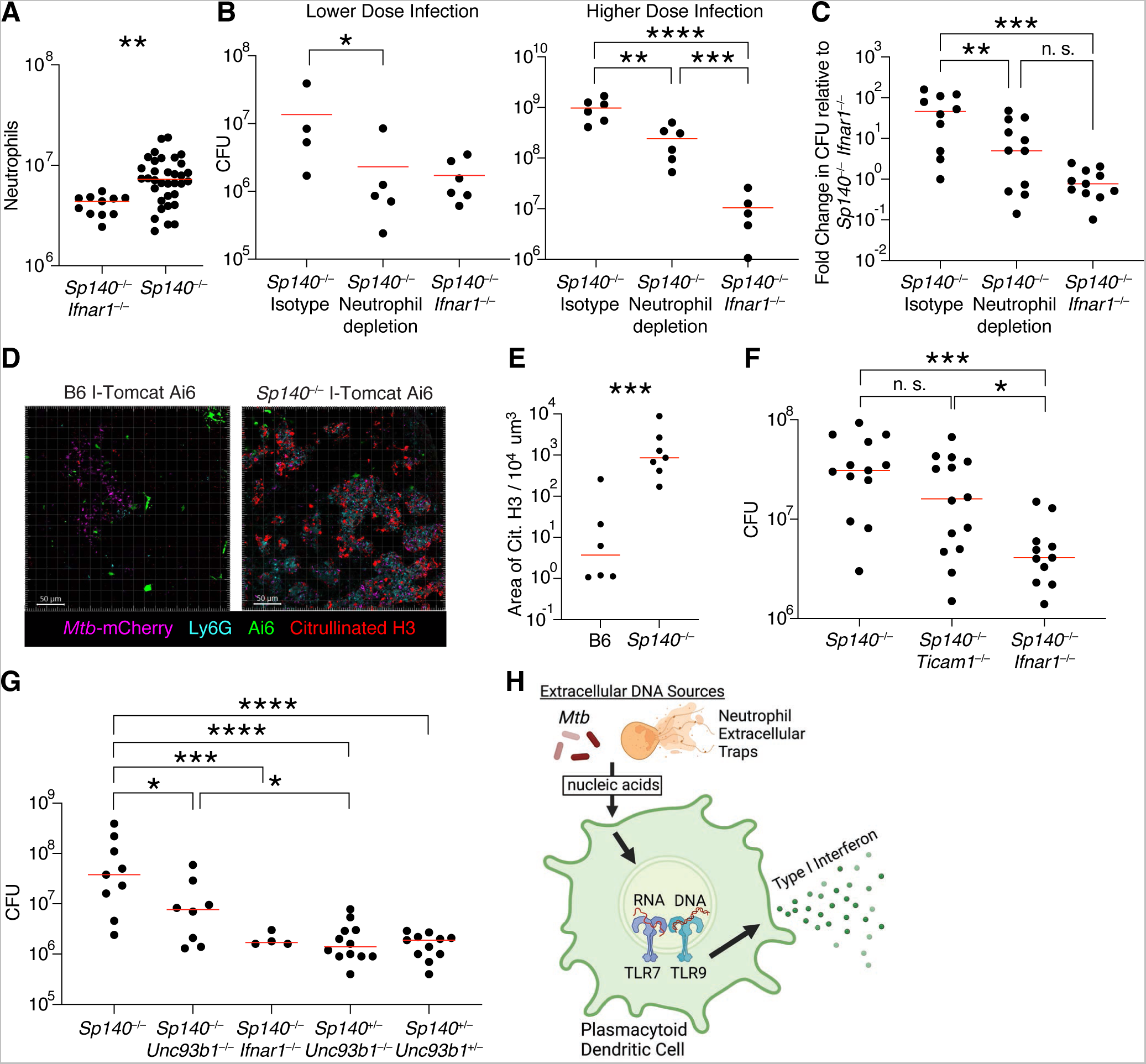
Role of neutrophils, neutrophil extracellular traps (NETs) and endosomal TLRs during type I IFN-driven *Mtb* pathogenesis. (**A**) Quantification of lung neutrophils in *Mtb*-infected *Sp140*^−/−^ (n = 34) and *Sp140*^−/−^ *Ifnar1*^−/−^ (n = 11) mice 25 days post-infection. (**B**) Bacterial burden after lower (n = 4-6) or higher dose (n = 5-6) infection and (**C**) combined normalized bacterial burden (n = 10-11) in *Sp140*^−/−^ mice that received isotype or neutrophil depleting anti-Ly6G clone 1A8 antibody from days 12 to 24 post-infection. (**D**) Representative images and (**E**) quantification of NET production based on citrullinated H3 staining in the diseased portions of I-Tomcat Ai6 and *Sp140*^−/−^ I-Tomcat Ai6 lungs. Sections were stained with citrullinated H3 (red), Ai6 (green), Ly6G (teal), and *Mtb*-expressed mCherry (magenta) (n = 6-7). (**F**) Lung bacterial burden in *Mtb*-infected *Sp140*^−/−^ (n = 13), *Sp140*^−/−^ *Ticam1*^−/−^ (*Ticam1* encodes TRIF; n = 14), and *Sp140*^−/−^ *Ifnar1*^−/−^ mice (n = 11). (**G**) Lung bacterial burden in *Mtb*-infected *Sp140*^−/−^ (n = 9), *Sp140*^−/−^ *Unc93b1*^−/−^ (n = 8), *Sp140*^−/−^ *Ifnar1*^−/−^ (n = 4), *Sp140*^+/−^ *Unc93b1*^−/−^ mice (n = 12), and *Sp140*^+/−^ *Unc93b1*^+/−^ mice (n = 11). (**H**) Model of potential extracellular DNA sources stimulating pDC production of type I IFNs through endosomal TLR signaling. The bars in (A), (B), (C), (E), (F), and (G) represent the median. Lungs were analyzed 25 days after *Mtb* infection. Statistical significance was calculated by one-way ANOVA with Tukey’s multiple comparison test for (C), (F) and (G), by one-tailed unpaired t test for (B), and by two-tailed unpaired t test for (A) and (E). *p < 0.05, **p < 0.01, ***p < 0.001, ****p < 0.0001.

### Neutrophils and IMs are the major sensors of type I IFNs during Mtb infection

Having identified pDCs, IMs, and monocytes as the major type I IFN producers during *Mtb* infection, we next sought to identify which cells responded to this type I IFN. As expected, IFNAR was uniformly expressed by all lung myeloid cells, and therefore not informative for identifying IFN responsive cells (**Fig. 7A**)^64^. However, comparing differentially expressed genes in B6 and *Sp140*^−/−^ neutrophils and IMs showed a clear induction of IFN stimulated genes in cells from *Sp140*^−/−^ animals (**Fig. 7B**). A major complication, however, is that many genes induced by type I IFN are also induced by IFN*γ*, and most existing studies do not distinguish the two. Therefore, we sought to develop type I IFN-specific and IFN*γ*-specific transcriptional signatures. RNA-sequencing analysis of human macrophages and mouse bone marrow-derived macrophages stimulated with IFN*γ*, IFN-β, TNF, transforming growth factor-β, or nothing were used to define cytokine induced genes (**Supplementary Fig. 12, 13**)^65^. Similar gene families were preferentially upregulated by type I or II IFNs in human and mouse macrophages and the resulting gene signatures for type I or II IFN responsiveness were highly unique to stimulation with IFN-β or IFN*γ* (**Supplementary Fig. 12**). We next applied the mouse gene signatures to the mouse lung myeloid scRNA-seq dataset. Strength of signature expression in naïve mice was used to define the threshold for classifying cells as an IFN*γ* or type I IFN responder (**Supplementary Fig. 13**). As expected, naïve mice had very few cells responding to either cytokine, while bystander and *Mtb*-infected cells responded strongly to type I and 2 IFNs (**Fig. 7C**, **Supplementary Fig. 13**). Interestingly, the type I IFN response was limited to IMs and neutrophils, even though monocytes and AMs were responsive to IFN*γ*. Potentially, differences in the localization of these cells could explain their differences in cytokine responsiveness. As expected, *Mtb*-infected neutrophils and IMs from *Sp140*^−/−^ mice exhibited a significant increase in type I IFN signaling relative to cells from B6 lungs (**Fig. 7D**). Consistent with considerable prior work demonstrating that type I IFNs impair responsiveness to IFN*γ*^16,32,33,66^, the *Sp140*^−/−^ mice harbored a distinct population of IMs that exhibited the signature of type I IFN-responsiveness but lacked the signature of IFN*γ* responsiveness (note the distinct population of blue IMs among the *Mtb*-infected cells in **Fig. 7C, 7D**). The reduction in IFN*γ* signaling in *Mtb*-infected IMs in *Sp140*^−/−^ mice correlated with reduced IFN*γ* receptor 1 expression on these cells in *Sp140*^−/−^ relative to B6 mice (**Supplementary Fig. 14**). This reduced IFN*γ* receptor 1 expression also correlated with increased induction of type I IFN stimulated genes and reduced expression of type II IFN stimulated genes (**Supplementary Fig. 14**). Since IFN*γ* is critical for control of intracellular *Mtb* replication, these results suggest that type I IFN might impair *Mtb* control at least in part by opening a niche of susceptible IMs that fail to respond to IFN*γ*.

**Figure 7.**
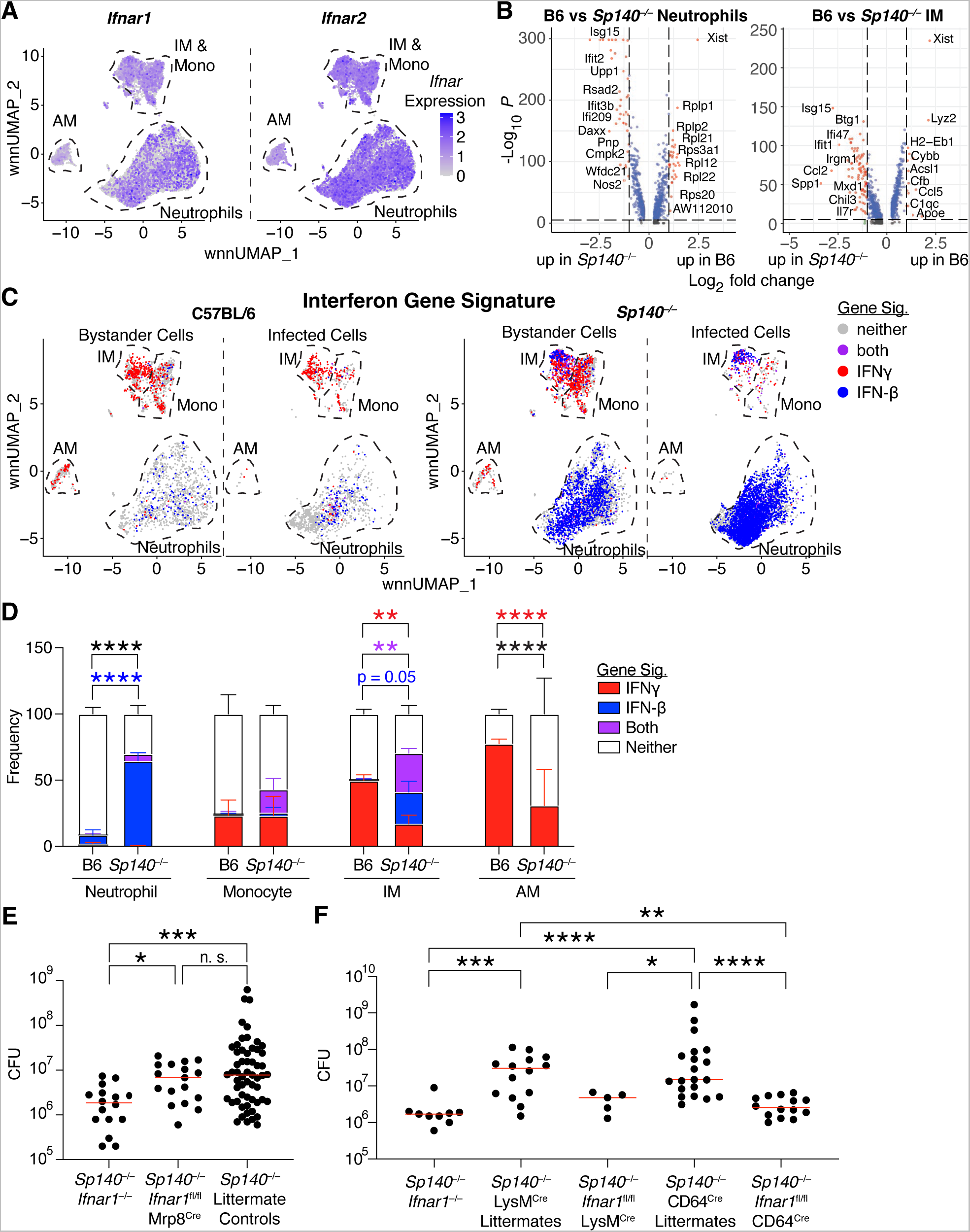
Macrophage recognition of type I IFN drives the *Mtb* susceptibility of *Sp140*^−/−^ mice. (**A**) wnnUMAP plot of *Ifnar1* and *Ifnar2* mRNA expression in innate immune cells from *Mtb*-infected lungs (B6 and *Sp140*^−/−^ combined). (**B**) Volcano plot of differentially expressed genes comparing B6 and *Sp140*^−/−^ neutrophils and IMs, with higher log fold change indicating greater expression in B6. (**C**) wnnUMAP plot of bystander, and *Mtb*-infected lung myeloid cells from B6 and *Sp140*^−/−^ mice classified by their responsiveness to IFNγ (red), type I IFN (blue), both (purple), or neither (grey). Signatures are defined in Supplementary Fig. 12 and applied for cell classification in Supplementary Fig. 13. (**D**) Graph of the frequency of neutrophils, monocytes, IMs, and AMs from *Mtb*-infected B6 (n = 3) and *Sp140*^−/−^ (n = 3) lungs that are responsive to IFN*γ* (red), type I IFN (blue), both (purple), or neither (white). (**E**) Lung bacterial burden in *Mtb*-infected *Sp140*^−/−^ *Ifnar1*^−/−^ (n = 16), *Sp140*^−/−^ *Ifnar1*^fl/fl^ Mrp8^cre^ (n = 18), and *Sp140*^−/−^ littermate control (n = 57) mice. (**F**) Lung bacterial burden in *Mtb*-infected *Sp140*^−/−^ *Ifnar1*^−/−^ (n = 9), *Sp140*^−/−^ LysM^cre^ littermate control (n = 14), *Sp140*^−/−^ *Ifnar1*^fl/fl^ LysM^cre^ (n = 5), *Sp140*^−/−^ CD64^cre^ littermate control (n = 20), and *Sp140*^−/−^ *Ifnar1*^fl/fl^ CD64^cre^ (n = 14) mice. The bars in (E) and (F) represent the median. Lungs were analyzed for the depicted experiments 24-26 days after *Mtb* infection. Pooled data from two-three independent experiments are shown. Statistical significance in (B) was calculated by non-parametric Wilcoxon rank sum test with Bonferroni correction, by one-way ANOVA with Tukey’s multiple comparison test for (E) and (F), and by two-way ANOVA with Tukey’s multiple comparisons test in (D). *p < 0.05, **p < 0.01, ***p < 0.001, ****p < 0.0001.

As neutrophils and IMs were the primary sensors of type I IFN, these are the cell types we tested to determine which cell population mediates the type I IFN-driven susceptibility of *Sp140*^−/−^ mice. Neutrophil-specific deficiency in type I IFN signaling was insufficient to rescue the *Mtb* susceptibility or the increase in lung neutrophils exhibited by *Sp140*^−/−^ mice (**Fig. 7E**, **Supplementary Fig. 15A**). Conversely, deletion of *Ifnar1* expression on myeloid cells with LysM^Cre^ *Ifnar1*^fl/fl^ mice or specifically in macrophages with CD64^Cre^ *Ifnar1*^fl/fl^ mice rescued bacterial control and reduced lung neutrophil numbers to the same extent as global *Ifnar1* deletion (**Fig. 7F**, **Supplementary Fig. 15B**). Taken together, these results are consistent with a model in which type I IFNs act on IMs to inhibit IFN*γ* signaling in these cells, thereby reducing their ability to restrict *Mtb* growth.

## Discussion

The dominant gene signature identified in humans with active tuberculosis disease is a type I IFN signature^7–10^. Type I IFNs are critical for effective anti-viral immunity^8^ but considerable data from humans and animal models indicate type I IFNs impair *Mtb* control^7,9,10,67^. For example, a range of viral co-infections or chemical interferon inducers exacerbate *Mtb* disease in mice^17–21^. Interestingly, although type I IFN and IFN*γ* (also called type II IFN) induce a highly overlapping set of target genes (**Supplementary Fig. 12**), the ability of type I IFNs to promote *Mtb* disease is not seen with IFN*γ*, which is instead potently protective against mycobacterial infections in mice^5^ and humans^68–70^. Indeed, type I IFNs can exacerbate bacterial infections by directly antagonizing IFN*γ* signaling^16,32,33,66^. The mechanism of antagonism is poorly understood but seems to be due at least in part to the downregulation of surface expression of the IFN*γ* receptor^32,33^. We see evidence for this type I IFN induced downregulation of IFN*γ* receptor during *Mtb* infection, as *Mtb*-harboring IMs express lower levels of IFN*γ* receptor (and IFNγ target genes) in *Sp140*^−/−^ mice, which produce a strong type I IFN response, relative to those in B6 mice, which produce a weak type I IFN response (**Supplementary Fig. 14**). The antagonism of IFN*γ* signaling by type I IFNs is conserved in humans, and has been shown to be relevant during *Mycobacterium leprae* infections^16^. Potentially, antagonism of IFN*γ* signaling underlies the tuberculosis-susceptibility of humans that exhibit a type I IFN signature, as humans with active *Mtb* exhibit reduced IFN*γ* receptor expression on their monocytes^15^. Type I IFNs can also impair IL-1 signaling, an additional pathway that is critical for *Mtb* control, through the induction of IL-1 receptor antagonist and eicosanoid imbalance^26,30,71,72^. Thus, the antagonism of protective IFNγ and IL-1 responses by type I interferons may be a key driver of progression to active tuberculosis. There is support in the literature for this hypothesis, as infants in their first year of life that acquire cytomegalovirus infection, a virus known to induce type I IFN, have a high risk of developing tuberculosis disease^11,12,73–75^. Additionally, influenza infection, which also induces type I IFN, correlates with an increased risk of death in pulmonary tuberculosis patients^13^. While the correlation between viral infection and exacerbated tuberculosis disease is tantalizing, a major limitation of studying human disease is the difficulty in establishing the underlying causal mechanisms that account for the correlation. Thus, we sought a genetically tractable model of *Mtb* infection in order to establish the cellular mechanisms by which type I IFN drives *Mtb* susceptibility.

The commonly used B6 mouse model does not exhibit a strong type I IFN response after *Mtb* infection^24,30,31^. Consistent with the modest type I IFN response of B6 mice, *Ifnar1* deletion on the B6 genetic background does not reliably impact survival or bacterial burdens in the lungs after *Mtb* infection^22–24,30^. Therefore, we sought a different mouse model that recapitulated two key aspects of human disease: the hyper type I IFN response, and the accompanying neutrophilic inflammation^7,76,77^. Previously, we identified B6.*Sst1*^S^ mice as a mouse model that exhibits type I IFN-driven susceptibility to *Mtb* infection^30^. We then demonstrated that the absence of *Sp140* in B6.*Sst1*^S^ mice explains their susceptibility to *Mtb*^31^. SP140 is a member of the Speckled Protein family of epigenetic readers and is widely expressed in leukocytes^78^. In humans, polymorphisms in *SP140* have been linked to Crohn’s disease^79,80^, chronic lymphocytic leukemia^81^, and multiple sclerosis^82^ by GWAS, but there is no known association between polymorphisms in *SP140* and susceptibility to *Mtb*. It has been suggested that macrophages in *Sp140*-deficient mice exhibit inherent defects in bacterial control causing them to be more susceptible to dextran sulfate sodium-induced colitis^39,40^. However, an inherent defect in bacterial control is not evident during *Mtb* infection, as *Sp140*^−/−^ mice lacking IFNAR restrict *Mtb* as well as B6 animals 25 days after infection^31^, a time point at which macrophages are critical for *Mtb* control^38^. This result suggests that the early susceptibility of *Sp140*^−/−^ mice is due to their strong type I IFN response rather an inherent defect in bacterial killing by macrophages. In addition to their hyper type I IFN response, *Sp140*^−/−^ mice also have more lung neutrophils after *Mtb* infection than *Sp140*^−/−^*Ifnar1*^−/−^ animals (**Fig. 1D, 7B, 7D**). Therefore, *Sp140*^−/−^ mice recapitulate the fundamental type I IFN and neutrophilic character of human active *Mtb* disease, and can serve as a platform for understanding the cellular mechanism of type I IFN-driven *Mtb* susceptibility.

*Sp140*^−/−^ mice provide a unique model of the aberrant type I IFN response, as they are on a pure B6 genetic background, and do not require repeated administration of TLR agonists, viral co-infection, or other perturbations of the innate immune system for type I IFN production^17,18,25,26,63^. Other groups have also modeled the type I IFN response by infecting B6 mice with a lineage 2 clinical *Mtb* strain, such as HN878^83,84^. However, *Ifnar1* deletion had no impact on survival or bacterial control at early time points in B6 mice infected with HN878, unlike *Sp140*^−/−^ mice infected with *Mtb* Erdman^31,85,86^. Thus, we believe the *Sp140*^−/−^ mouse model uniquely recapitulates the hyper type I IFN response exhibited by humans, and provides a tool to study the mechanistic basis of the aberrant type I IFN response.

We sought to use the *Sp140*^−/−^ mice to address two key questions: (1) what cells produce type I IFN during *Mtb* infection? and (2) which cells respond to the type I IFN to mediate *Mtb* disease? Flow cytometry and imaging of I-Tomcat Ai6 *Ifnb1* reporter mice identified IMs and pDCs as the major IFN-β producers during *Mtb* infection. Imaging provided insight into why these cells expressed type I IFN, as the frequency of type I IFN-expressing macrophages was enriched relative to CD4^+^ T cells in diseased tissue but not in healthy tissue. This result suggests that proximity to *Mtb* dictates access to activating signals required to induce IFN-β expression by macrophages. However, most IFN-β expressing IMs were not infected with *Mtb*, and most infected IMs were not *Ifnb1* or reporter positive, indicating direct infection is insufficient and may not be the main driver of type I IFN expression *in vivo*. *In vitro* studies have shown that bone marrow-derived macrophages infected with *Mtb* induce type I IFN via the cGAS-STING pathway. However, this pathway does not seem to be of great importance *in vivo*^30,87–90^. Instead, our results suggest that uninfected bystander cells may be the primary producers of type I IFN during *Mtb* infection. Consistent with this hypothesis, we found that mice lacking *Unc93b1*, a chaperone required for TLRs that sense extracellular nucleic acids, exhibit enhanced control of *Mtb*. Exogenous nucleic acids sensed by uninfected bystander cells may thus be an important pathway for type I IFN induction during *Mtb* infection *in vivo*. Of note, pDCs, which we found to be important type I IFN producers during *Mtb* infection, are robust producers of type I IFN after TLR7/9 sensing of exogenous nucleic acids^58,91,92^. Imaging demonstrated that B6 and *Sp140*^−/−^ macrophages expressed IFN-β at similar frequencies in diseased tissue. This result appears to contrast with flow cytometry analysis of whole lungs, which demonstrates a higher frequency of IMs expressing IFN-β in *Sp140*^−/−^ mice relative to B6 mice. The apparent discrepancy between the flow cytometry and imaging data is likely explained by the fact that *Sp140*^−/−^ mice have considerably more diseased tissue than B6 mice. Additionally, the Ai6 signal from I-Tomcat Ai6 mice identifies cells that have expressed IFN-β, but does not indicate the level of IFN-β expression in these cells. Thus, even if macrophages are driven to express IFN-β at similar frequencies in B6 and *Sp140*^−/−^ mice, the macrophages in *Sp140*^−/−^ mice could produce more IFN-β per cell, as seen in the scRNA-seq data (**Fig. 3C**).

Expression of type I IFNs by pDCs during *Mtb* infection was particularly noteworthy as limited work exists on the effect of pDCs on *Mtb* control. pDCs are believed to be important for control of viral infections, with their production of type I IFN during the early phases of viral infections significantly contributing to viral control^56,57,61^. During bacterial infections, the potential for pDCs to impact control has become appreciated more recently, with pDCs demonstrating a protective function against *Citrobacter rodentium*, *Chlamydia pneumoniae*, and *Klebsiella pneumoniae*^93–96^. However, few studies have examined the contribution of pDCs during *Mtb* infection, and these studies have not yet led to a clear understanding. The number of pDCs in the blood was reduced in *Mtb*-infected humans, but lung pDC numbers or function were not assessed^97,98^. In non-human primates, there was a correlation between active pulmonary *Mtb* and the influx of pDCs and IFN-responsive macrophages into the lungs of the rhesus macaques^51^. Another group also identified pDCs in tuberculosis granulomas of a different non-human primate, cynomolgus macaques, but the frequency of pDCs among all cells in the granulomas did not correlate with bacterial burden of the granuloma^99^. A major issue in the studies using NHPs or humans is the difficulty in depleting or otherwise functionally assessing the role of pDCs. To address this limitation, we generated *Sp140*^−/−^ pDC-DTR mice to study the contribution of pDCs to bacterial control. These mice are the most specific tool currently available for deletion of pDCs during *Mtb* infection^59,61^ and demonstrated that pDCs (or pDC-like cells) contribute significantly to the susceptibility of *Sp140*^−/−^ mice. Additionally, we identified pDCs in the lymphocytic cuff surrounding *Mtb* granulomas in human lungs and lymph nodes. Together, these results suggest that disease progression driven by pDC production of type I IFN drives disease in mice, and is likely conserved across non-human primates and humans.

While pDC depletion rescued *Sp140*^−/−^ mice, it had no impact on bacterial burden in B6 animals. This result was expected given that very few myeloid cells in B6 mice expressed a type I IFN signature and *Ifnar1* deficiency also has only modest effects in the B6 background^24,30^. The specific lack of an effect of pDC depletion in *Sp140*-sufficient mice also suggests that the role of pDCs in exacerbating *Mtb* infection is related to their production of type I IFNs instead of an IFN-independent function. As pDCs are present in B6 and *Sp140*^−/−^ mice, we speculated that the difference in pDC type I IFN production in these mouse strains could be due in part to differences in the availability of activating ligands. As seen in another *Mtb* susceptible mouse model, and in *Mtb*-infected human lungs^63^, *Sp140*^−/−^ mice had a significant enrichment in NET production compared to B6 mice. Potentially, these NETs, which are DNA-rich products of neutrophils, act as ligands for TLR9 on the pDCs, as described in mouse and human autoimmunity^34,35,62,100^. In support of this hypothesis, neutrophil depletion rescued the susceptibility of *Sp140*^−/−^ mice, as did deletion of *Unc93b1* which is required for the function of endosomal TLRs, including TLR9 which senses DNA. Given that NET formation was also identified in *Mtb* granulomas in human lung sections^63^, it is possible that pDC sensing of NETs contributes to the type I IFN response detected in humans with active *Mtb* disease.

Having defined the cells producing type I IFNs *in vivo* after *Mtb* infection, we then sought to identify which cells responded to the type I IFN. To do this, we had to first develop transcriptional signatures that distinguish the response to type I IFN from the closely related response to IFN*γ*. Applying these signatures to our scRNA-seq data, we identified neutrophils and IMs as type I IFN sensors. While both cell populations harbor *Mtb*, it is possible that type I IFN driven susceptibility to *Mtb* is caused by signaling in only one of these populations. In a GM-CSF blockade model of type I IFN-driven *Mtb* susceptibility, neutrophil-specific deletion of *Ifnar1* rescued bacterial control^63^. We employed the same genetic strategy as this prior report, but were unable to detect any rescue of *Sp140*^−/−^ mice when neutrophils lacked *Ifnar1*. This was surprising as we saw that *Sp140*^−/−^ neutrophils expressed a type I IFN gene signature by scRNA-seq. However, GM-CSF is critical for maintaining lung alveolar macrophages and the responsiveness of lung monocytes and macrophages to infections, including *Mtb* infection^101–104^. Therefore, it is possible that impairing lung macrophages by GM-CSF blockade shifted the impact of type I IFN on *Mtb* control from macrophages to neutrophils. In support of this idea, we also identified IMs as a type I IFN-sensing population, and deletion of *Ifnar1* on macrophages rescued *Sp140*^−/−^ mouse bacterial control. Our scRNA-seq dataset only contains myeloid cells, therefore we would miss the contribution of type I IFN signaling in other cell types. However, it is likely that type I IFN largely acts through myeloid cells to drive *Mtb* susceptibility, as macrophage-specific deletion of *Ifnar1* rescued *Mtb* control to a similar extent as global *Ifnar1* deficiency. These results suggest that during a normal (GM-CSF sufficient) response, type I IFN signaling in macrophages reduces their ability to restrict *Mtb*. The mechanism by which type I IFNs impair *Mtb* clearance by IMs remains unknown. However, an observation to emerge from our studies was that infected IMs in *Sp140*^−/−^ mice exhibit a reduced transcriptional signature of responsiveness to IFNγ, and instead primarily exhibited a type I IFN signature, consistent with the exacerbated type I IFN response in these mice. The transcriptional response of *Sp140*^−/−^ IMs contrasted dramatically with that of infected IMs in B6 mice, which control *Mtb* and which exhibited a uniform signature of responsiveness to IFNγ (**Figure 7**). Given the essential role of IFNγ in control of *Mtb* in mice and humans, our results suggest that one detrimental effect of type I IFNs may potentially be via inhibition of IFN*γ* signaling in the macrophages^15,32,33^.

Taken together, our results identify the cell types that produce and respond to type I IFNs during IFN-driven tuberculosis disease. We propose that type I IFNs impair responsiveness to IFN*γ*, leading to an initial loss of bacterial control. Bacterial replication then leads to neutrophil influx and NET production within the diseased tissue. We propose that DNA-rich NETs are sensed by pDCs, likely via endosomal TLRs, causing them to produce type I IFN, which further antagonizes IFN*γ* signaling and reduces the IMs ability to restrict *Mtb* growth. Given the correlations between our results and findings in rhesus macaques and humans with active *Mtb*, we believe that our proposed mechanism of type I IFN driven loss of *Mtb* control is conserved across species. These findings open the door for the development of therapies targeting NET production or pDC function as host-directed strategies for treating active *Mtb* infection.

## Limitations of the Study

While we see a strong correlation between NET formation and type I IFN-driven *Mtb* susceptibility, the current study does not directly test the contribution of NETs in this response. Additionally, the present study provides data suggesting that type I IFN signaling correlates with a loss of type II IFN signaling in IMs during *Mtb* infection, but does not directly examine whether *Mtb*-harboring IMs in *Sp140*^−/−^ mice are unable to control *Mtb* infection because of a lack of response to IFN*γ*. We also limited our studies to ~25 days post-infection, which is an early time point for *Mtb* infection. It is possible that the cellular sources and targets of type I IFN shift to other cell types at later time points in the infection.

## Acknowledgments

We thank members of the Vance, Barton, Stanley, and Cox laboratories for advice and discussions; M. Colonna for advice on pDC depletion; B. Malissen, Y. Belkaid, and A. Lacy-Hulbert for mouse lines; P. Dietzen, R. Chavez, and J. Morales for technical assistance; A. Valeros, H. Nolla, and K. Heydari of the UC Berkeley Cancer Research Laboratory Flow Cytometry facility for assistance with flow cytometry; H. Aaron and F. Ives of the UC Berkeley Cancer Research Laboratory Molecular Imaging Center (RRID:SCR_017852) for assistance with confocal microscopy (supported by the Helen Wills Neuroscience Institute). Model figures were created with BioRender.com. D.I.K. was supported by an NIH Postdoctoral Fellowship (F32HL158250). R.E.V. is an Investigator of the Howard Hughes Medical Institute and is funded by NIH grants AI075039, AI063302 and AI155634.

## Author Contributions

Conceptualization: D.I.K. and R.E.V. Investigation: D.I.K., O.V.L., S.A.F, C.L., J.M.P., A.M., and E.B. Data analysis: D.I.K. Methodology: J.G., K.C.W., and D.B.S. Resources: D.B.S. Writing: D.I.K. and R.E.V. with input from all co-authors. Funding acquisition: R.E.V. Supervision: R.E.V., D.B.S., D.L.J., and B.D.B.

## STAR Methods

### RESOURCE AVAILABILITY

#### Lead contact

Further information and requests for resources and reagents should be directed to and will be fulfilled by the lead contact, Russell Vance.

#### Materials availability

Materials used in this study will be provided upon request and available upon publication.

#### Data and code availability

- Raw and processed bulk RNA- and single cell RNA-sequencing data is deposited in the NCBI Gene Expression Omnibus: GSE216023, GSE232827, GSE232922. This paper also analyzes existing, publicly available data, for which the accession numbers are listed in the key resources table.
- Code for bulk RNA- and scRNA-sequencing analysis is available on Github: https://github.com/dmitrikotov/Sp140-Type-I-Inteferon.
- Any additional information required to reanalyze the data reported in this paper is available from the lead contact upon request.

### EXPERIMENTAL MODEL AND STUDY PARTICIPANT DETAILS

#### Animals

Mice were maintained under specific pathogen-free conditions and housed at 23°C with a 12 hour light-dark cycle in accordance with the regulatory standards of the University of California Berkeley Institutional Animal Care and Use Committee. All mice were sex- and age-matched and were 6-12 weeks old at the start of infections. Male and female mice were used in all experiments. Littermate controls were used when possible, as indicated in the figure legends. B6, B6.129S2-Ifnar1^tm1Agt^/Mmjax (*Ifnar1*^−/−^)^105^, B6.Cg-Gt(ROSA)26Sor^tm6^(CAG–ZsGreen1)^Hze^/J (Ai6)^52^, B6(Cg)-Ifnar1^tm1.1Ees^/J (*Ifnar1*^fl^)^106^, B6.Cg-Tg(S100A8-cre,-EGFP)1Ilw/J (Mrp8^Cre^)^107^, B6.129P2-Lyz2^tm1(cre)Ifo^/J (LysM^Cre^)^108^, and C57BL/6J-*Ticam1^Lps^*^2^/J (*Ticam1*^−/−^)^109^ mice were purchased from Jackson Laboratories. C57BL/6N-*Unc93b1^tm^*^1*(KOMP)Vlcg*^/Mmucd (*Unc93b1*^−/−^) mice were obtained from the Mutant Mouse Resource and Research Center (MMRRC) at the University of California, Davis, was donated to the MMRRC by the KOMP repository at University of California, Davis, originated from David Valenzuela of Regeneron Pharmaceuticals^110^, and were provided by G. Barton at the University of California, Berkeley. *Ifnb1*-Tomato-Cre-pA Terminator (I-Tomcat) mice were a gift from D. Stetson at the University of Washington and a manuscript describing these mice in detail is currently in preparation. B6-Fcgr1^tm2Ciphe^ (CD64^Cre^)^111^ mice were a gift from B. Malissen at Centre d’Immunologie de Marseille Luminy and were provided by Y. Belkaid at the National Institutes of Health. B6-Tg(CLEC4C-HBEGF)956Cln/J (pDC-DTR)^61^ mice were provided by A. Lacy-Hulbert at the Benaroya Research Institute. *Sp140*^−/−^ mice were previously generated in-house^31^. *Sp140*^−/−^*Ifnar1*^−/−^ mice were generated by crossing *Sp140*^−/−^ mice with *Ifnar1*^−/−^ mice in-house. I-Tomcat Ai6 mice were generated by crossing I-Tomcat mice with Ai6 mice, while *Sp140*^−/−^ I-Tomcat Ai6 mice were the result of crossing I-Tomcat mice with *Sp140*^−/−^ and Ai6 mice in-house. *Sp140*^−/−^ mice were crossed in-house with pDC-DTR mice to generate *Sp140*^−/−^ pDC-DTR mice. *Sp140*^−/−^ *Ifnar1*^fl^ LysM^cre^, *Sp140*^−/−^ *Ifnar1*^fl^ Mrp8^Cre^, and *Sp140*^−/−^ *Ifnar1*^fl^ CD64^Cre^ mice were generated by crossing *Sp140*^−/−^ mice with *Ifnar1*^fl^ and LysM^Cre^ or Mrp8^Cre^ or CD64^Cre^ mice in-house.

#### Bacterial strains

*Mtb* strain Erdman was a gift of S. A. Stanley. Frozen aliquoted stocks were produced after passing the strain *in vivo* to ensure virulence. *Mtb* expressing Wasabi (*Mtb*-Wasabi) and *Mtb*-mCherry were generated using *Mtb* that had been passaged 2 or fewer times *in vitro*. For these fluorescent strains, *Mtb* was grown in Middlebrook 7H9 liquid medium supplemented with 10% albumin-dextrose-saline, 0.4% glycerol, and 0.05% Tween-80 for 5 days at 37°C. The cells were pelleted and washed in 10% glycerol to remove salt. The bacteria were then electroporated with 1 µg DNA using a 2 mm electroporation cuvette and the following settings: 2500 volts, 1000 Ohms, 25 µF. The pTEC15 plasmid (a gift from Lalita Ramakrishnan; Addgene plasmid # 30174), which expresses Wasabi under the control of the Mycobacterium Strong Promoter, was electroporated into *Mtb* to generate *Mtb*-Wasabi^36^. The pMSP12::mCherry plasmid (a gift from Lalita Ramakrishnan; Addgene plasmid # 30167), which expresses mCherry under the control of the Mycobacterium Strong Promoter, was electroporated into *Mtb* to generate *Mtb*-mCherry. Following electroporation, bacteria were grown on 7H11 plates supplemented with 10% oleic acid, albumin, dextrose, and catalase, 0.5% glycerol, and either 200 µg / mL Hygromycin for *Mtb*-Wasabi or 50 µg / mL Kanamycin for *Mtb*-mCherry for 3-4 weeks at 37°C. Individual colonies where then propagated in 10 mL inkwell flask cultures using 7H9 medium supplemented with 10% albumin-dextrose-saline, 0.4% glycerol, 0.05% Tween-80, and either Hygromycin for *Mtb*-Wasabi or Kanamycin for *Mtb*-mCherry for 7 days at 37°C. The inkwell cultures were expanded into a 100 mL culture using the same 7H9 supplemented media with antibiotics and cultured for 4-5 days at 37°C. Once the bacteria were in log phase, the culture was filtered with a 5 µm syringe filter and frozen in 1 mL aliquots in 10% glycerol.

### METHOD DETAILS

#### *Mtb* infections

For infection, a frozen aliquot of *Mtb*-Wasabi or *Mtb*-mCherry was diluted in distilled H_2_O, and 9 mL of diluted culture was loaded into the nebulizer of a inhalation exposure system (Glas-Col, Terre Haute, IN) to deliver ~20-100 bacteria per mouse as determined by measuring CFU in lungs 1 day post-infection.

#### Tissue Processing for CFU and Flow cytometry

Mice were harvested at various days post-infection (as described in figure legends) to measure CFUs by plating and innate immune populations by flow cytometry. All lung lobes were harvested into a gentleMACS C tube (Miltenyi Biotec) containing 3 mL of RPMI media with 70 µg / mL of Liberase TM (Roche) and 30 µg / mL of Dnase I (Roche). Samples were processed into chunks using the lung_01 setting on the gentleMACS (Miltenyi Biotec) and incubated for 30 minutes at 37°C. Tissue was then homogenized into a single cell suspension by running the samples on the lung_02 setting on the gentleMACS. The digestion was quenched by adding 2 mL of PBS with 20% Newborn Calf Serum (Thermo Fisher Scientific) and filtered through 70 µm SmartStrainers (Miltenyi Biotec).

For measuring plasmacytoid dendritic cell numbers, spleens were harvested into a 12 well plate with 1 mL of PBS with 2% Newborn Calf Serum and 0.05% sodium azide in each well. The spleens were sandwiched between 100 uM mesh filters and mashed into a single cell suspension with the back of a syringe plunger. The single cell suspensions were filtered through 70 µm SmartStrainers (Miltenyi Biotec).

#### Measuring Bacterial Burden

To measure CFU, 50 µL was taken from each single cell suspension and then serially diluted in phosphate-buffered saline (PBS) with 0.05% Tween-80. Serial dilutions were plated on 7H11 plates supplemented with 10% oleic acid, albumin, dextrose, and catalase and 0.5% glycerol. Colonies were counted after 3 weeks.

#### Flow Cytometry

For flow cytometry, lung single cell suspensions were pelleted and resuspended in 500 µL of PBS with 2% Newborn Calf Serum and 0.05% Sodium azide and 100-150 µL were stained with antibodies for analysis. Spleen single cell suspensions were pelleted and resuspended in 5 mL of PBS with 2% Newborn Calf Serum and 0.05% Sodium azide, of which 50 µL were stained with antibodies. Single cell suspensions were stained for 45 minutes to an hour at room temperature with the following antibodies: TruStain FcX PLUS (S17011E, BioLegend), BUV496- labeled CD45 (30-F11, BD Biosciences), APC-labeled CD64 (X54-5/7.1, BioLegend), BV480-labeled B220 (RA3-6B2, BD Biosciences), BV480-labeled CD90.2 (53-2.1, BD Biosciences), APC-Fire 750-labeled Ly6G (1A8, BioLegend), BUV395-labeled CD11b (M1/70, BD Biosciences), BUV737-labeled CD11c (HL3, BD Biosciences), APC-R700-labeled Siglec F (E50-2440, BD Biosciences), PE-labeled MerTK (DS5MMER, Thermo Fisher Scientific), Super Bright 645-labeled MHC II (M5/114.15.2, Thermo Fisher Scientific), BV421-labeled PD-L1 (MIH5, BD Biosciences), BV711-labeled Ly6C (HK1.4, BioLegend), PE-labeled IFNAR-1 (MAR1-5A3, BioLegend), PE-Cy7-labeled MerTK (DS5MMER, Thermo Fisher Scientific), APC-eFluor 780-labeled CD11b (M1/70, Thermo Fisher Scientific), BUV395-labeled CCRL2 (BZ2E3, BD Biosciences), BUV563-labeled Ly6G (1A8, BD Biosciences), Percp-Cy5.5-labeled B220 (RA3-6B2, BioLegend), BV421-labeled Siglec H (440c, BD Biosciences), BV480-labeled CD19 (1D3, BD Biosciences), BV605-labeled MHC II (M5/114.15.2, BioLegend), BV785-labeled Ly6C (HK1.4, BioLegend), BV605-labeled CD4 (GK1.5, BioLegend), BUV805-labeled CD8ɑ (53-6.7, BD Biosciences), and PE-Cy7-labeled PDCA-1 (eBio927, Thermo Fisher Scientific). All samples also received fixable viability dye (Ghost Dye Violet 510; Tonbo Biosciences), Super Bright Complete Staining Buffer (Thermo Fisher Scientific), and True-Stain Monocyte Blocker (BioLegend) at the same time as the antibodies. Stained samples were fixed with cytofix/cytoperm (BD biosciences) for 20 minutes at room temperature before samples were removed from the BSL3. For intracellular staining, the fixed samples were stained with the following antibodies: PE-Cy7-labeled CD63 (NVG-2, BioLegend), Percp-eFluor 710-labeled iNOS (CXNFT, Thermo Fisher Scientific), BV785-labeled CD206 (C068C2, BioLegend). Cell numbers were calculated by adding fluorescent AccuCheck Counting Beads (Invitrogen) to each sample. Cells were then analyzed on a Fortessa (BD Biosciences) or an Aurora (Cytek) flow cytometer. Data were analyzed with Flowjo version 10 (BD Biosciences).

#### Immune cell depletion

Injections for cell depletions were started 12 days after *Mtb* infection and continued every other day until the mice were harvested 25 days post-infection. Antibody depletion of pDCs was performed by administering 200 µg of anti-BST2 (927, BioXCell) or rat IgG2b isotype control antibody (LTF-2, BioXCell) in 200 µL PBS via intraperitoneal injection^59^. Genetic depletion involved administering 100 ng diphtheria toxin (Millipore Sigma) in 100 µL PBS via intraperitoneal injection into *Sp140*^−/−^ pDC-DTR mice and littermate controls^61^. Antibody depletion of neutrophils was performed by administering 200 µg of anti-Ly6G (1A8, BioXCell) or rat IgG2a isotype control antibody (2A3, BioXCell) in 200 µL PBS via intraperitoneal injection.

#### Immunostaining human lymph nodes and lungs

Human lung and lymph node samples were acquired from the surgical pathology archives of Emory University Hospital with appropriate institutional approval. 8 lung samples and 8 lymph node samples were analyzed. Each sample had been previously culture verified for *Mycobacterium tuberculosis* infection. The samples were formalin-fixed and paraffin-embedded. Sections were cut and stained with anti-CD123 (6h6, Thermo Fisher Scientific), anti-CD303 (124B3.13, Dendritics), or hematoxylin and eosin^112,113^. Primary antibodies were detected by immunoperoxidase staining with the LSAB+ System and a standard DAB reaction following manufacturer’s instructions (DakoCytomation). Sections were counterstained with hematoxlyin prior to mounting and microscopy. pDCs were assessed in multiple 400× fields for each section to calculate the frequency of samples containing pDCs and the clustering of pDCs within each sample, defined as either singe cells, loose clusters of 5-20 cells, or tight clusters of more than 20 cells.

#### Confocal microscopy

Confocal Microscopy was performed using a Zeiss LSM 880 laser scanning confocal microscope (Zeiss) equipped with two photomultiplier detectors, a 34-channel GaASP spectral detector system, and a 2-channel AiryScan detector as well as 405, 458, 488, 514, 561, 594, and 633 lasers. 20 µm paraformaldehyde fixed lung sections from *Mtb*-mCherry infected I-Tomcat Ai6 and *Sp140*^−/−^ I-Tomcat Ai6 mice were stained at 4°C overnight with BV421-labeled SIRP⍺ (P84, BD Biosciences), Pacific Blue–labeled B220 (RA3-6B2, BioLegend), eF506-labeled CD4 (RM4-5, BioLegend), and AF647-labeled Ly6G (1A8, BioLegend). Sections detecting the presence of Neutrophil extracellular traps (NETs) were stained with BV421-labeled Ly6G (1A8, BioLegend) and rabbit polyclonal anti-citrullinated histone-H3 (citrulline R2, R8, R17; Abcam), stained with AF488 donkey anti-rabbit secondary (Poly4064, BioLegend). Stained sections were inspected with a 5× air objective to find representative lesions and distal sites and then imaged using a 63× oil immersion objective lens with a numerical aperture of 1.4. For each infected lung, one *Mtb*-heavy lesion image and one distal site image was taken consisting of 20 µm z-stacks acquired at a 1.5 µm step size. For representative NET images, 4×4 tiled images were captured without a z-stack. Additionally, the Zeiss LSM 880 microscope was used to image single color-stained Ultracomp eBeads Plus (Thermo Fisher Scientific) for generating a compensation matrix.

#### Image processing and histo-cytometry analysis

Image analysis was performed using Chrysalis software^55^. Briefly, a compensation matrix was generated by automatic image-based spectral measurements on single color-stained controls in ImageJ by using Generate Compensation Matrix script. This compensation matrix was used to perform linear unmixing on three-dimensional images with Chrysalis. Chrysalis was also used for further image processing, including rescaling data and generating new channels by performing mathematical operations using existing channels. For histo-cytometry analysis, Imaris 9.9.1 (Bitplane) was used for surface creation to digitally identify cells in images based on protein expression^54^. Statistics for the identified cells were exported from Imaris and then imported into FlowJo version 10 (BD Biosciences) for quantitative image analysis.

#### Bulk RNA-seq sample preparation and analysis

RNA-seq of *Mtb*-infected *Sp140*-sufficient and -deficient mouse lungs genetically depleted of pDCs was performed on 20% of each lung single cell suspension, prepared as described for CFU and flow cytometry analysis. Single cell suspensions were preserved in Trizol LS (Thermo Fisher Scientific) and stored at −80°C. The samples were thawed at room temperature for 5 minutes, then 200 µL of chloroform (Thermo Fisher Scientific) was added per 0.75 mL of Trizol LS to each sample and the samples were removed from the BSL3. Samples were centrifuged in Phasemaker tubes (Thermo Fisher Scientific) to isolate total RNA, which was then purified following the RNeasy Micro (Qiagen) protocol starting at the ethanol addition step. Library preparation, sequencing, and read alignment to the mouse genome was performed by Azenta Life Sciences. Libraries were prepared from total RNA using an Illumina kit for Poly(A) selection. Samples were sequenced on an Illumina HiSeq sequencer with paired 150 bp reads to a depth of 20-30 million reads per sample. Sequence reads were trimmed of adapter sequences and low quality nucleotides with Trimmomatic v.0.36^114^ and then mapped to the Mus musculus GRCm38 reference genome with STAR aligner v.2.5.2b^115^. The raw counts were used as input for DESeq2^116^ analysis of differential gene expression.

RNA-seq of cytokine stimulated bone marrow-derived macrophages was performed by differentiating bone marrow from B6 mice in DMEM supplemented with 10% FBS, PenStrep, Glutamin, HEPES, and M-CSF for 7 days then reseeding the cells in a 6-well plate and incubating for 5 days at 37°C and 5% CO_2_. The macrophages were left untreated or stimulated with 10 ng / mL of IFN-β (BioLegend), IFN*γ* (Abcam), TNF (Peprotech), or transforming growth factor-β (BioLegend) for 6 hours. Cells were lysed with TRK lysis buffer (Omega Bio-Tek) and 2-mercaptoethanol (Thermo Fisher Scientific) followed by total RNA isolation using the E.Z.N.A Total RNA Kit I (Omega Bio-Tek). The library preparation, sequencing, and read alignment to the mouse genome was performed by Azenta Life Sciences as described for the *Mtb*-infected mouse lung samples. Raw counts were used as input for analysis with DESeq2^116^.

#### Sorting Immune Cells for scRNA-seq analysis

3 B6 and 3 *Sp140*^−/−^ mice were infected with less than 100 CFU of *Mtb*-Wasabi bacteria. Lungs from infected animals as well as 2 naïve control mice per genotype were harvested 25 days post-infection and processed as described for CFU and flow cytometry analysis. Single cell suspensions were resuspended in PBS with 2% Newborn Calf Serum and stained with TruStain FcX PLUS (S17011E, BioLegend), APC-labeled Ly6G (1A8, BioLegend), APC-labeled CD64 (X54-5/7.1, BioLegend), and TotalSeq-A-labeled Ly6G (1A8, BioLegend) anti-mouse antibodies on ice for 30 minutes. Myeloid cells were then magnetically enriched using an EasySep APC Positive Selection Kit II (StemCell Technologies) and MojoSort Magnets (BioLegend). All enriched samples were stained with the following panel of TotalSeq-A-labeled anti-mouse antibodies to detect protein expression in the scRNA-seq dataset: Ly6C (HK1.4, BioLegend), CD44 (IM7, BioLegend), CD169 (3D6.112, BioLegend), CD274 (MIH6, BioLegend), Siglec F (S17007L, BioLegend), CSF1R (AFS98, BioLegend), CD11b (M1/70, BioLegend), CD86 (GL-1, BioLegend), MHC II (M5/114.15.2, BioLegend), and CX3CR1 (SA011F11, BioLegend). Each sample was also stained with a unique anti-mouse TotalSeq-A Hashtag antibody (1-6; BioLegend) to allow up to 6 populations to be multiplexed in a single lane on a 10X Genomics Chromium Next GEM Chip^117^. Enriched cells from infected mice were also stained with PE-labeled B220 (RA3-6B2, Tonbo), PE-labeled CD90.2 (30-H12, Tonbo), and BV785-labeled CD45.2 (104, BioLegend) anti-mouse antibodies. Enriched cells from naïve mice were stained with Pacific Blue-labeled B220 (RA3-6B2, BioLegend), Pacific Blue-labeled CD90.2 (53-2.1, BioLegend), PE-labeled F4/80 (BM8, Thermo Fisher Scientific), and BV785-labeled CD45.2 (104, BioLegend) anti-mouse antibodies. All post-enrichment antibody staining was performed on ice for 45 minutes in the presence of True-Stain Monocyte Blocker (BioLegend). Following staining, cells were resuspended in PBS with 2% Newborn Calf Serum and Sytox Blue Dead Cell Stain (Thermo Fisher Scientific). Cells were sort purified using a 100 µm microfluidic sorting chip in a 4 laser SH-800 cell sorter (Sony) on the purity setting. The isolated populations from infected lungs were *Mtb*-infected cells and bystander myeloid cells. Macrophages and a mixture of neutrophils and monocytes were isolated from the naïve lungs and the macrophages were combined with neutrophil/monocyte mixture at a 1:2 ratio for better macrophage representation in the resulting dataset.

#### Single cell RNA: Library generation and sequencing

The scRNA-sequencing libraries were generated using the v3.1 chemistry Chromium Single Cell 3’ Reagent Kit (10X Genomics) largely following the manufacturer protocol with the following minor modifications. Cells were loaded into 3 different lanes on a Chromium Next GEM Chip. Lane 1 was loaded with *Mtb*-infected cells from all 3 B6 and 3 *Sp140*^−/−^ lungs. Lane 2 was loaded with bystander myeloid cells from the 3 infected B6 lungs as well as the myeloid cell mixture from the 2 naïve B6 lungs. Lane 3 was loaded with bystander myeloid cells from the 3 infected *Sp140*^−/−^ lungs as well as the myeloid cell mixture from the 2 naïve *Sp140*^−/−^ lungs. All 3 lanes of the Chromium Next GEM Chip were super-loaded with 29000 cells with a target of 14800 single cells per lane, as hashtag barcoding allows for a lower effective multiplet rate due to the ability to identify most of the multiplets (https://satijalab.org/costpercell/)^117^. 0.5 U/µL RNaseOUT Recombinant Ribonuclease Inhibitor (Invitrogen) was added to single cell RT master mix during the loading step and 1 µL of ADT and HTO additive primers (0.2 µM stock) were added during the cDNA amplification, as recommended by the CITE-seq and Cell Hashing Protocol (https://cite-seq.com/protocols/)^42^. Following cDNA, ADT, and HTO purification, samples were decontaminated by 2 rounds of centrifugation through 0.2 µM filter microcentrifuge tubes and then removed from the BSL3. Library preparations were completed outside of the BSL3 following the 10X Genomics protocol for the cDNA and the CITE-seq and Cell Hashing Protocol for the ADT and HTO libraries. Quality control of the libraries was performed with a Fragment Analyzer (Agilent). The mRNA, ADT, and HTO libraries were pooled at the following proportions: 85% mRNA, 9% ADT, and 6% HTO. Libraries were sequenced on a NovaSeq 6000 (Illumina) using two lanes of a S1 flow cell and the following cycles read 1 (28 cycles), i7 index (10 cycles), i5 index (10 cycles), read 2 (90 cycles).

#### ScRNA-seq: data processing

Raw sequencing reads for the mRNA libraries were processed into raw count matrices with CellRanger version 4.0.0 (10X Genomics). The ADT and HTO libraries were processed into raw count matrices with CITE-Seq-Count version 1.4.3 (https://hoohm.github.io/CITE-seq-Count/)^118^. The raw counts for mRNA, ADT, and HTO were analyzed in R^119^ via the RStudio integrated development environment with Seurat v4.1.1^43^ using default settings for normalizing the data, finding variable features, and scaling the data. HTO demultiplexing was performed with the HTODemux function. Data was filtered to only include single cells with between 200 and 4500 genes and less than 5% mitochondrial reads. The resulting datasets were integrated together using 30 dimensions for the FindIntegrationAnchors function and 30 dimensions for the IntegrateData function. The data was then scaled and analyzed by PCA with 30 principal components followed by UMAP analysis with 30 dimensions. Clustering was performed by using 30 dimensions with the FindNeighbors function and a resolution of 0.8 for the FindClusters function.

To improve resolution for clustering innate immune cells, weighted nearest neighbor analysis was used to combine the protein data (ADTs) and the mRNA data when clustering cells. For this analysis, variable ADT features were identified and then normalized using centered log ratio transformation and a margin of 2. The normalized ADT data was then scaled and analyzed by PCA. The ADT and mRNA data was then combined with the FindMultiModalNeighbors function using 30 dimensions for the mRNA and 10 for the protein. The resulting dataset was analyzed by UMAP and clusters were identified with the FindClusters function using algorithm 3 and a resolution of 1.5^44^. Tidyverse^120^, EnhancedVolcano^121^, and various Seurat functions were used for plotting the scRNA-seq data.

#### Type I IFN and IFN*γ* Gene Signature Analysis

For generating the type I IFN and IFN*γ* gene signatures, we utilized a published RNA-seq dataset (GEO: GSE20251) of primary human macrophages that were unstimulated or stimulated with 10 ng/mL of TNF, IFN*γ*, IFN-β, transforming growth factor-β, or other ligands for 24 hours and then processed for RNA-sequencing^65^. We also generated an RNA-seq dataset mouse bone marrow-derived macrophages left unstimulated or stimulated with 10 ng/mL of TNF, IFN*γ*, IFN-β, or transforming growth factor-β for 6 hours. For both datasets, counts were normalized by DESeq2’s median of ratios method and biological replicates were averaged to generate average normalized counts for each cytokine condition. The average normalized counts were used to calculate a max to second-max ratio for the 4 cytokine conditions to determine gene specificity. Next, gene expression was compared across all stimulation conditions and filtered to only include genes that were induced relative to the untreated cells by a log_2_ fold change of 1 by IFN*γ* or IFN-β and to exclude genes induced by a log_2_ fold change of 1.5 by TNF or transforming growth factor-β. Using this filtered gene list, a ratio was calculated of the log_2_ fold change following IFN*γ* stimulation to the log_2_ fold change following IFN-β stimulation. For the human dataset, IFN-β specific genes were defined as those having a fold change ratio < 0, a log_2_ fold change upon IFN-β stimulation > 1, an average normalized count following IFN-β stimulation > 2000 and a max to second-max ratio > 2.5. Human IFN*γ* specific genes were defined as those with a fold change ratio > 1.5, a log_2_ fold change upon IFN*γ* stimulation > 2, an average normalized count following IFN*γ* stimulation > 1000 and a max to second-max ratio > 2.5. Mouse macrophage IFN-β specific genes were defined as those with a fold change ratio < 0.66, a log_2_ fold change upon IFN-β stimulation > 4, an average normalized count following IFN-β stimulation > 4000 and a max to second-max ratio > 3. Mouse IFN*γ* specific genes were defined as those that had a fold change ratio > 1.5, a log_2_ fold change upon IFN*γ* stimulation > 2, an average normalized count following IFN*γ* stimulation > 500 and a max to second-max ratio > 3.

Human IFN-β and IFN*γ* gene signatures were validated by examining their induction following human macrophage stimulation with a variety of cytokines, such as IL-4, IL-6 and IL-10, using the published RNA-seq dataset (GEO: GSE20251). The specificity of the mouse macrophage IFN*γ* signature was validated by stimulating bone marrow-derived macrophages for 6 hours with 10 ng/mL of IFN*γ*, IFN-β, or nothing and then examining CXCL9 expression by flow cytometric analysis of intracellular staining with PE-labeled anti-mouse CXCL9 antibody (MIG-2F5.5; BioLegend). The mouse gene signatures were then used to score cells in the mouse myeloid scRNA-seq dataset based on their gene expression with the UCell R package^122^.

### QUANTIFICATION AND STATISTICAL ANALYSIS

Group sizes were informed by the results of preliminary experiments and power calculations. The number of animals in each figure is indicated in the legends as n = x mice per group. Statistical significance was determined using Prism (GraphPad) software for unpaired one-tailed or two-tailed Student t test when comparing two populations, one-way or two-way ANOVA tests with Tukey’s or Sidak’s multiple comparisons test when comparing multiple groups. Prism (GraphPad) was also used to calculate linear correlations and R^2^. R was used to calculate statistical significance for bulk RNA- and scRNA-sequencing datasets using the Wald test with multiple testing correction by the Benjamini and Hochberg method and the Wilcoxon Rank-Sum test with Bonferroni correction, respectively. *p < 0.05, **p < 0.01, ***p < 0.001, ****p < 0.0001, n.s. = not significant. See figure legends for more information on statistical tests.

## Supplementary Figure Legends

**Supplementary Figure 1.**
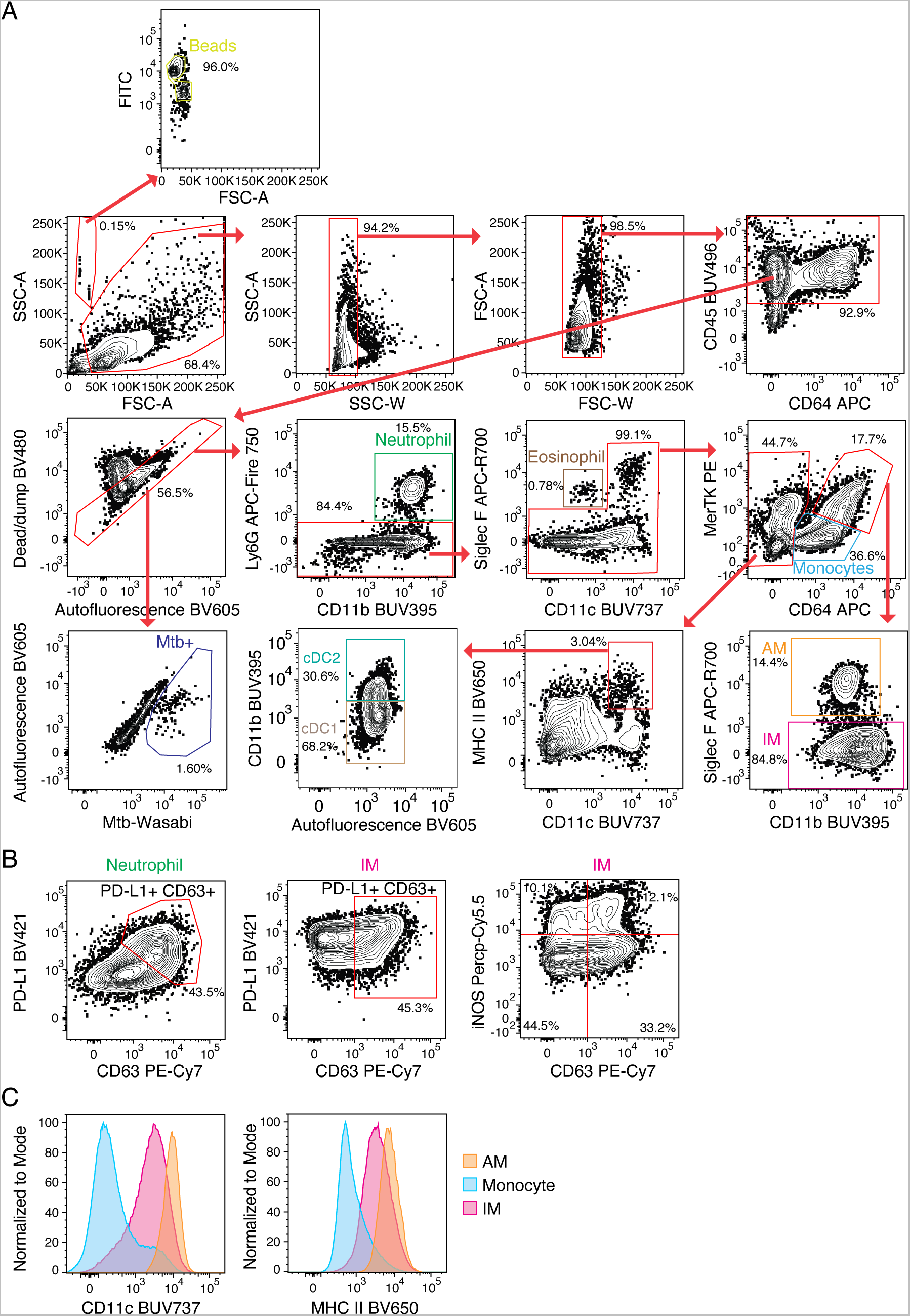
Representative flow cytometry gating strategy to identify innate immune cell populations in *Mtb*-infected lungs. (**A**) Gating strategy for identifying neutrophils, eosinophils, monocytes, dendritic cells (DCs), alveolar macrophages (AMs), interstitial macrophages (IMs), conventional type 1 dendritic cells (cDC1), conventional type 2 dendritic cells (cDC2) and *Mtb*-infected cells. Very few *Mtb*^+^ cells were detected among the lymphoid and non-hematopoietic-derived cells. (**B**) Representative identification of activated (CD63^+^ PD-L1^+^ or iNOS^+^) neutrophils and interstitial macrophages in *Mtb*-infected lungs. (**C**) Histogram comparing CD11c and MHC II expression on AMs (orange), monocytes (blue), and IMs (red).

**Supplementary Figure 2.**
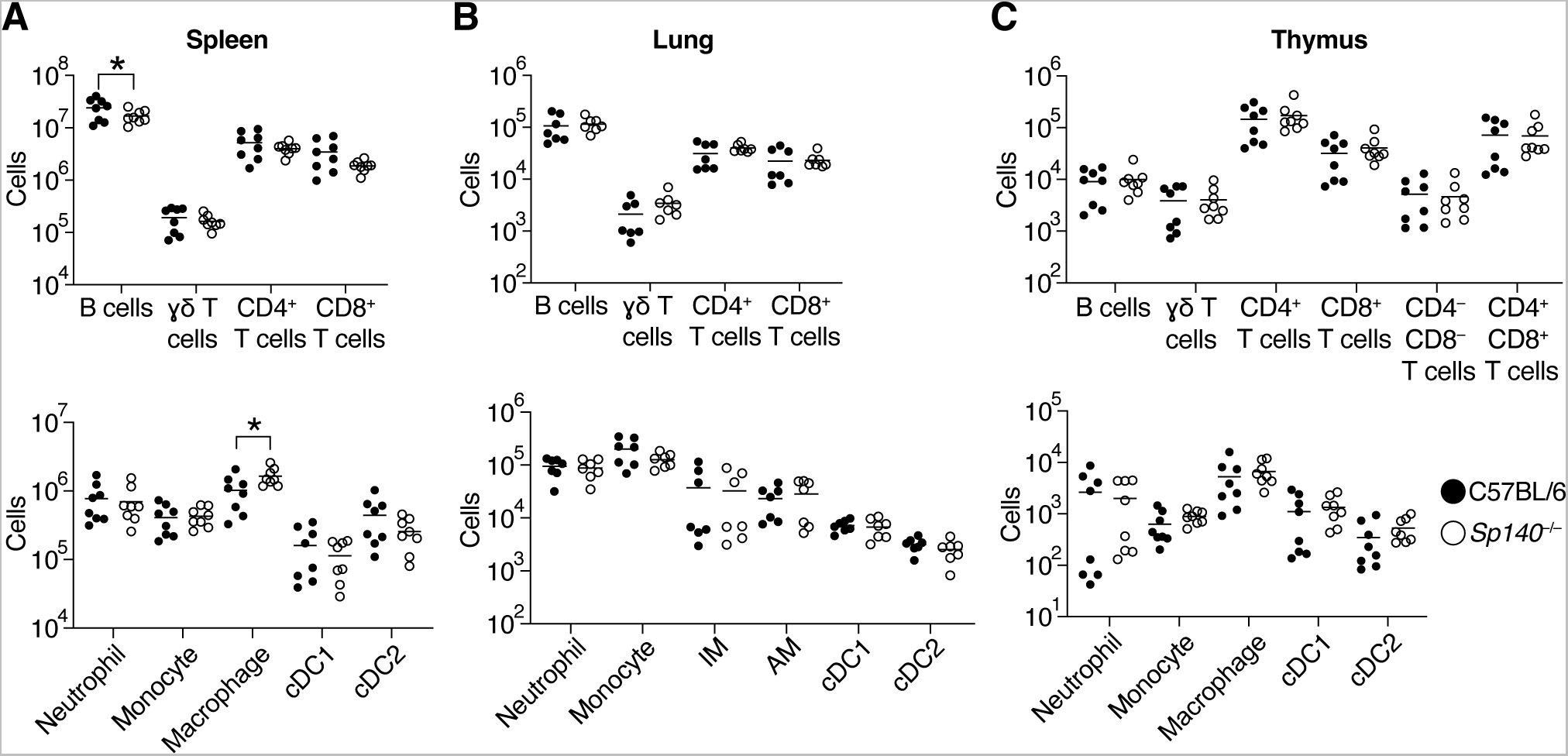
Naïve *Sp140*^−/−^ mice exhibit minimally altered immune cell numbers by flow cytometry. Comparison of innate and adaptive immune cell numbers between B6 (n = 7; closed circles) and *Sp140*^−/−^ (n = 8; open circles) (**A**) spleen, (**B**) lung, (**C**) and thymus. The bars in (A), (B), and (C) represent the median. Pooled data from two independent experiments are shown in (A), (B), and (C). Statistical significance was calculated by multiple unpaired t tests in (A), (B), and (C).

**Supplementary Figure 3.**
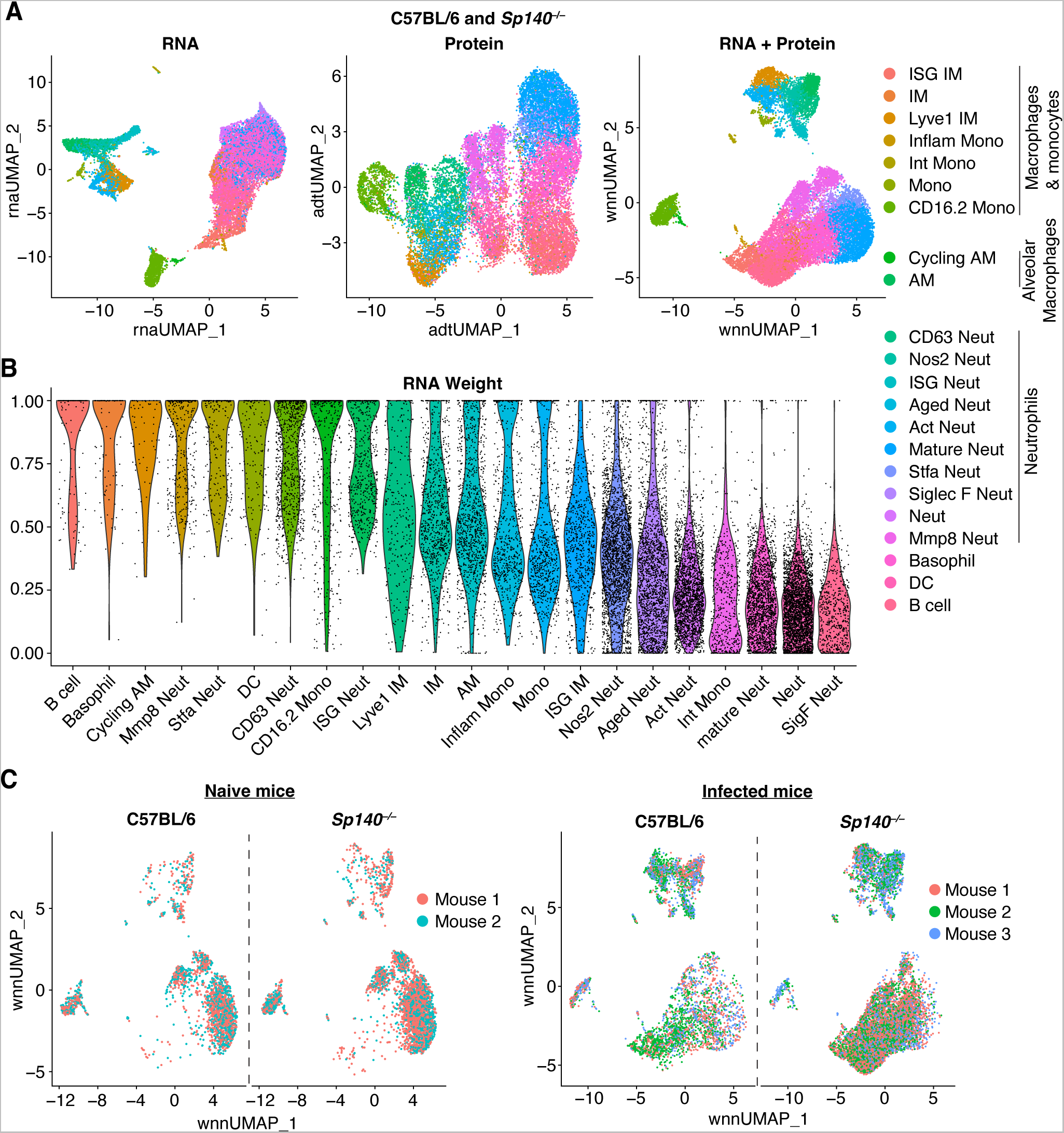
The scRNA-seq dataset is a CITE-seq experiment that represents multiple mice and combines mRNA and protein expression. (**A**) UMAP plots depicting cell clustering by mRNA expression, protein expression, or combined mRNA and protein expression (B6 and *Sp140*^−/−^ combined). (**B**) Violin plot of the RNA weight of each cluster, thereby depicting clusters identified based primarily on mRNA (closer to RNA weight = 1) or protein (closer to RNA weight = 0; B6 and *Sp140*^−/−^ combined). (**C**) wnnUMAP plot of the biological replicates from naïve and infected B6 and *Sp140*^−/−^ mice. For each genotype, n = 3 for the infected lung samples and n = 2 for the naïve lung samples.

**Supplementary Figure 4.**
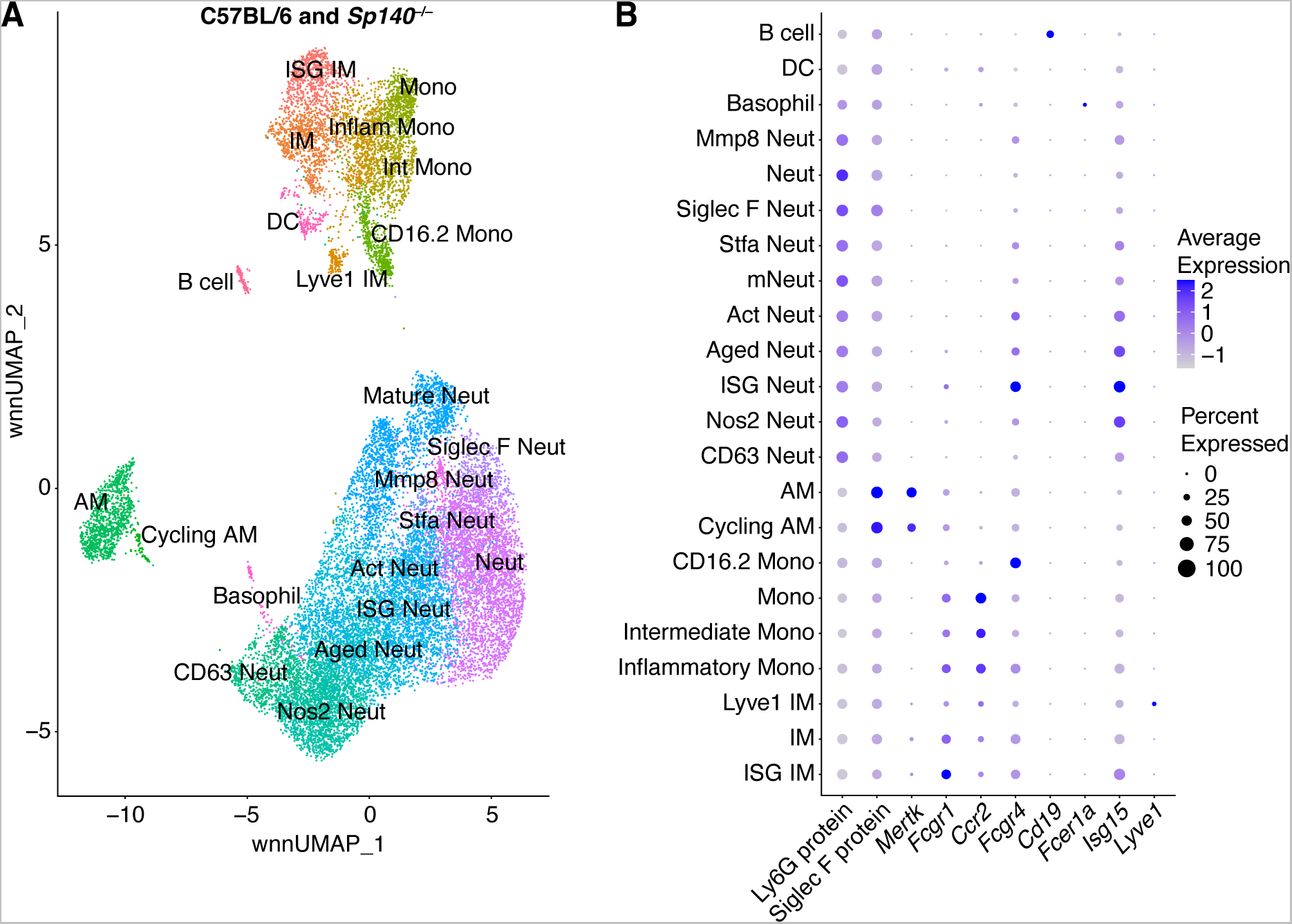
Lineage classification of myeloid cells from naïve and *Mtb*-infected B6 and *Sp140*^−/−^ mice. (**A**) wnnUMAP plot of myeloid cells from naïve and *Mtb*-infected mice (B6 and *Sp140*^−/−^ combined) with each cluster labeled. (**B**) Protein and mRNA expression of lineage defining markers used for cluster classification. The data represents a total of 10 mice (n = 3 for the infected lung samples and n = 2 for the naïve lung samples from B6 and *Sp140*^−/−^ mice).

**Supplementary Figure 5.**
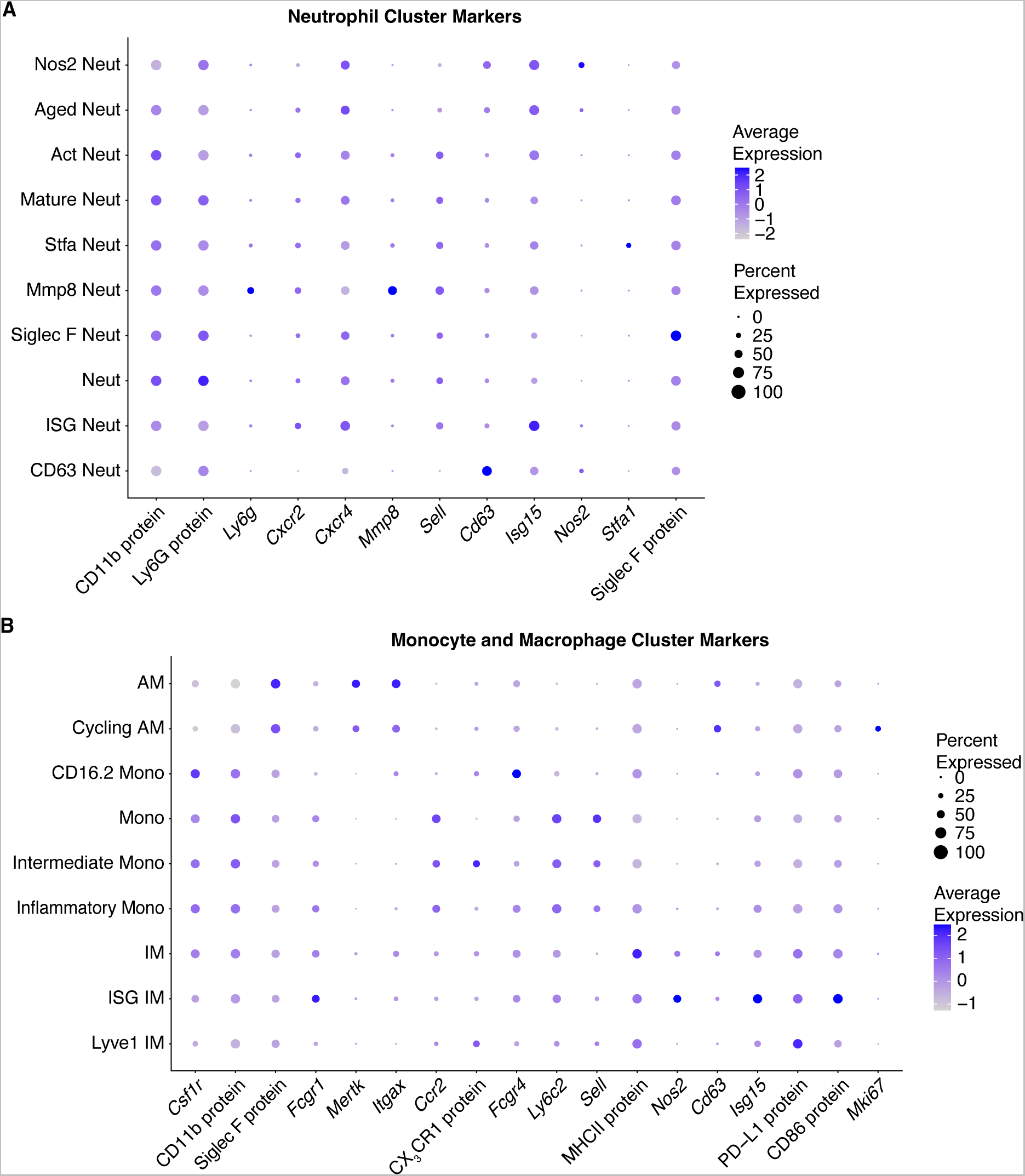
The expression of protein and mRNA used to annotate the myeloid cell clusters in the scRNA-seq dataset. Markers used for annotating (**A**) neutrophil and (**B**) monocyte and macrophage clusters based on maturity, activation status, and unique gene expression. The data represents a total of 10 mice (n = 3 for the infected lung samples and n = 2 for the naïve lung samples from B6 and *Sp140*^−/−^ mice).

**Supplementary Figure 6.**
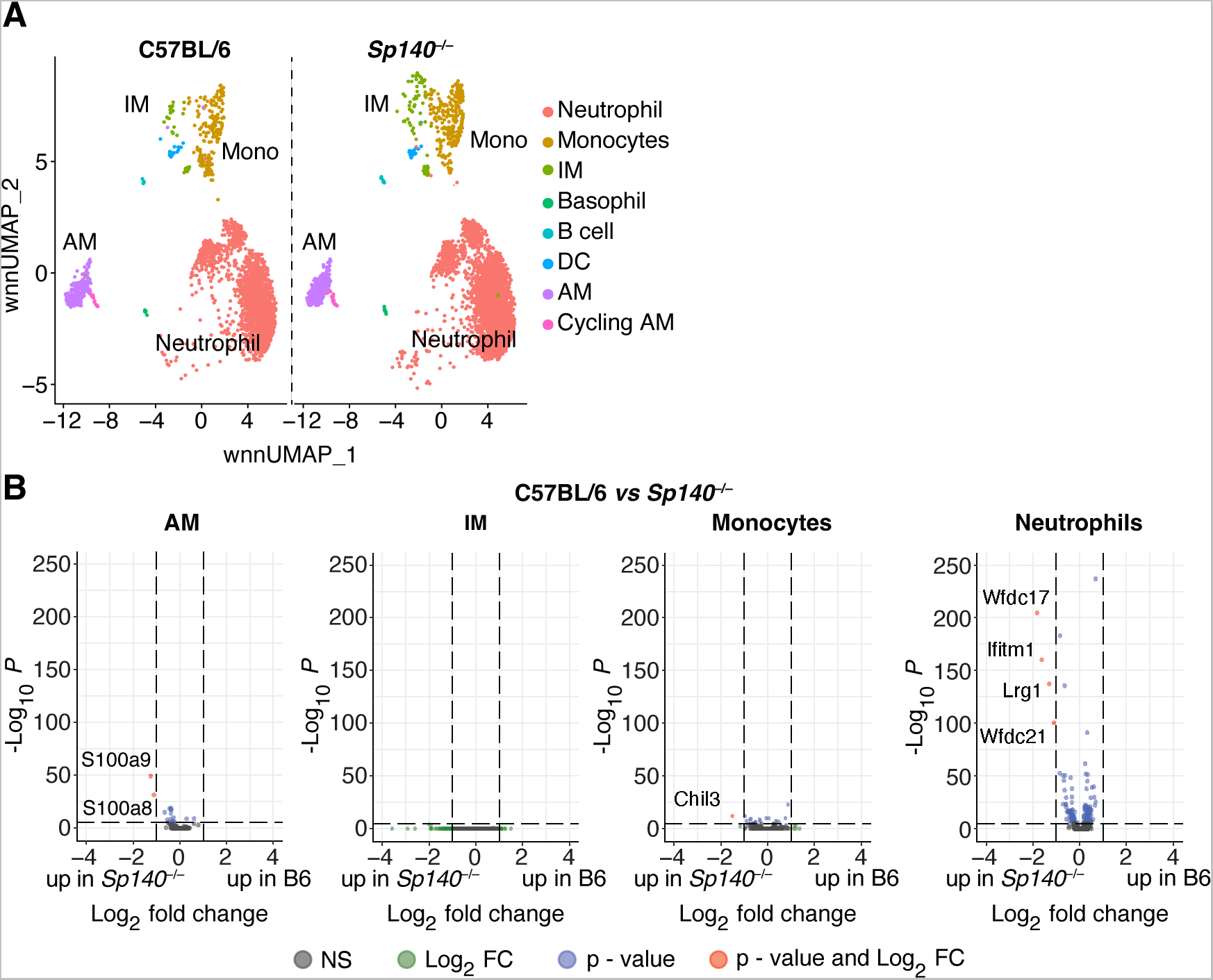
Naïve *Sp140*^−/−^ mice do not exhibit increased inflammation by scRNA-seq. (**A**) wnnUMAP plot of myeloid cells from naïve lungs of B6 (n = 2) and *Sp140*^−/−^ (n = 2) mice. (**B**) Volcano plot of the differentially expressed genes between B6 and *Sp140*^−/−^ alveolar macrophages (AMs), interstitial macrophages (IMs), monocytes, and neutrophils. Greater fold change indicates higher expression in B6 relative to *Sp140*^−/−^. Statistical significance in (B) was calculated with the Wilcoxon Rank-Sum test with Bonferroni correction.

**Supplementary Figure 7.**
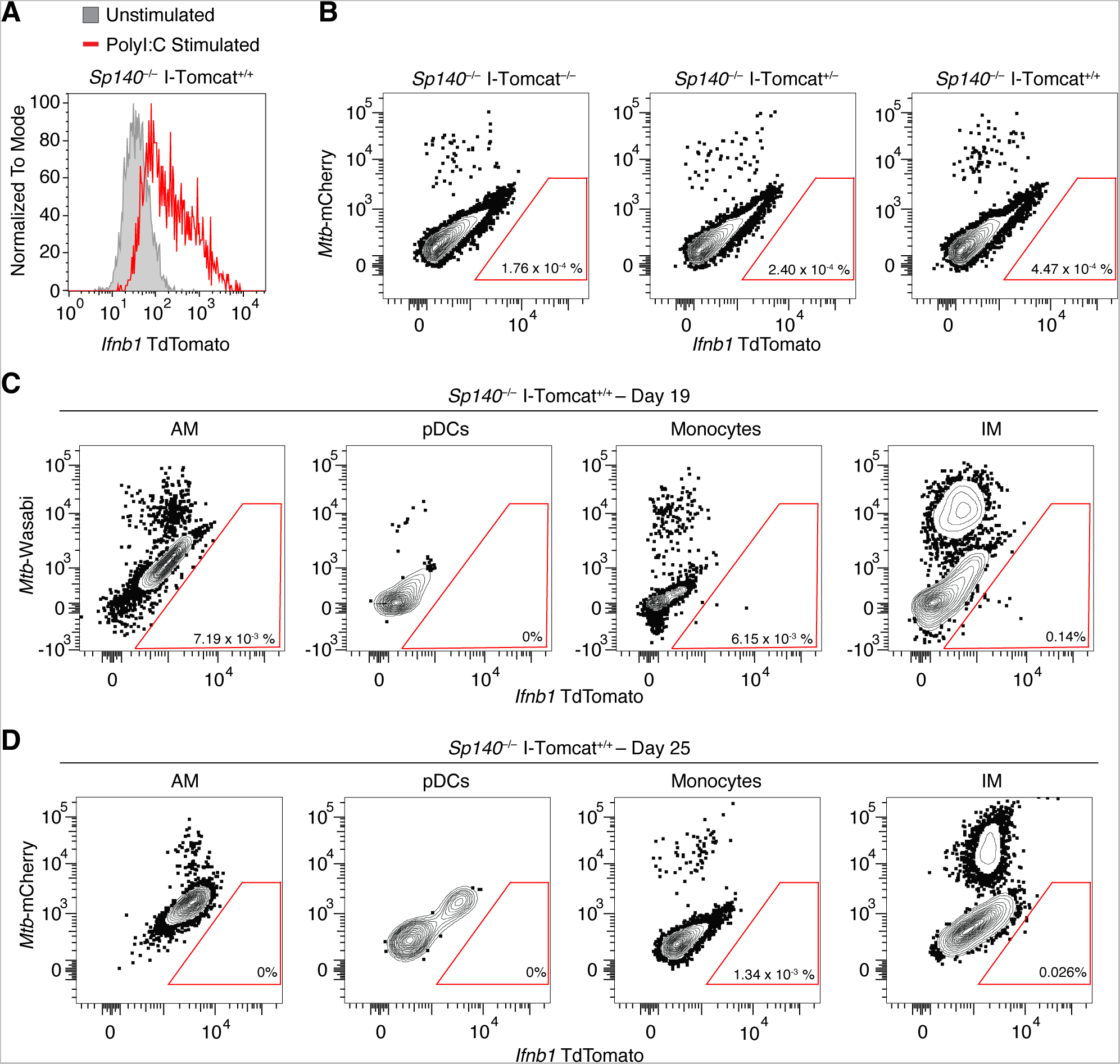
I-Tomcat bone marrow-derived macrophages express TdTomato following *in vitro* poly I:C stimulation, but the TdTomato signal is insufficient to detect IFN-β producing cells *in vivo* during *Mtb* infection. (**A**) Representative flow cytometry histogram of TdTomato expression by I-Tomcat bone marrow-derived macrophages that were unstimulated (grey) or stimulated with poly I:C (red line). (**B**) Representative flow cytometry plot of TdTomato expression in immune cells in *Mtb*-infected lungs of *Sp140*^−/−^ I-Tomcat^−/−^, *Sp140*^−/−^ I-Tomcat^+/−^, and *Sp140*^−/−^ I-Tomcat^+/+^ mice 25 days after infection. (**C**) Representative flow cytometry plot of TdTomato expression in AM, pDCs, monocytes, and IM from *Mtb*-infected lungs of *Sp140*^−/−^ I-Tomcat^+/+^ mice 19 days post-infection or (**D**) 25 days post-infection.

**Supplementary Figure 8.**
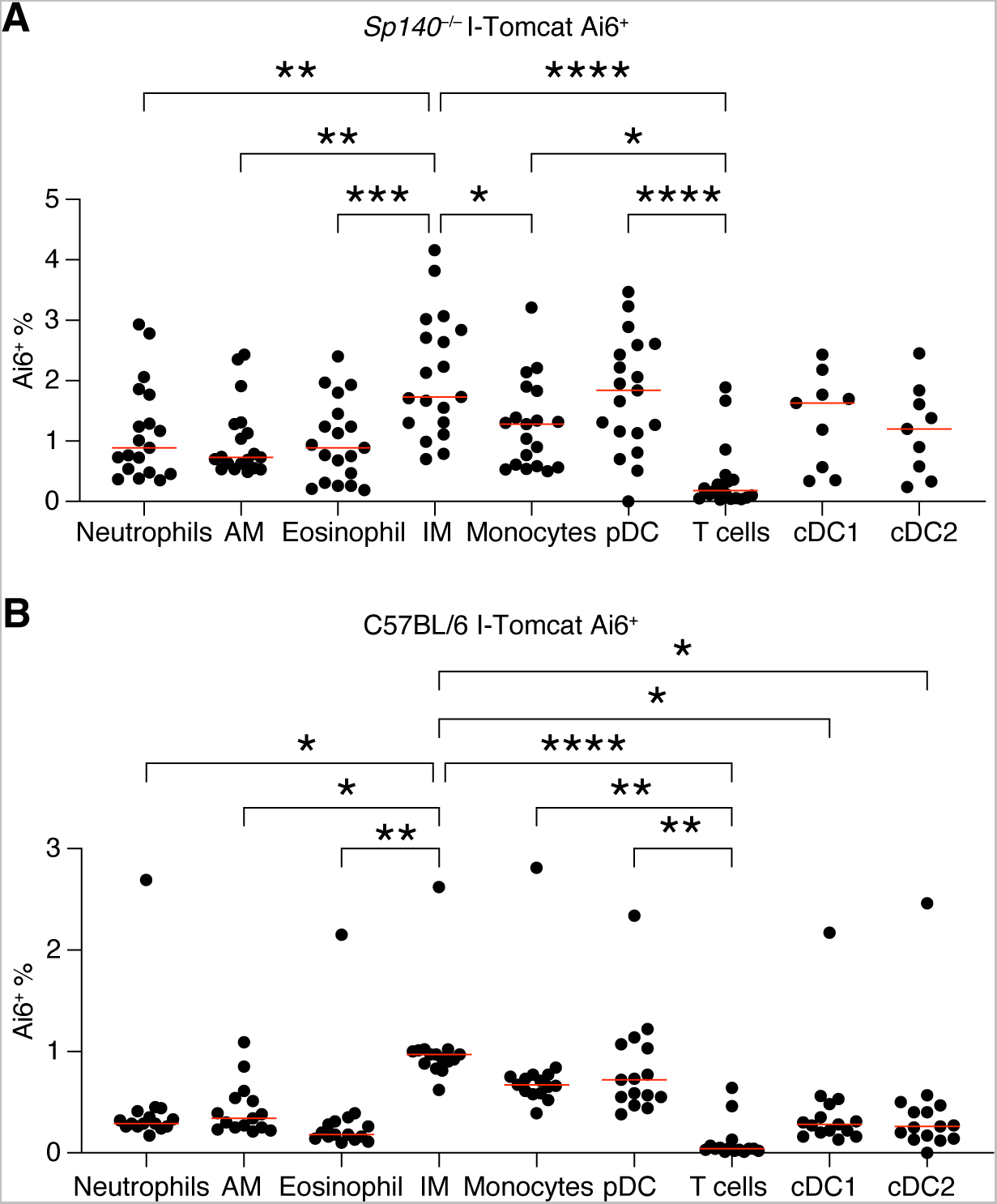
pDC, IMs, and monocytes are the major type I interferon producing cells following *Mtb* infection in B6 and *Sp140*^−/−^ mice. Frequency of Ai6 expression by immune cell type in the lungs of *Mtb*-infected (**A**) *Sp140*^−/−^ I-Tomcat Ai6 and (**B**) I-Tomcat Ai6 mice. The bars in (A) and (B) represent the median. Lungs were analyzed 25 days after *Mtb* infection. Pooled data from two independent experiments. Statistical significance was calculated by one-way ANOVA with Tukey’s multiple comparison test. *p < 0.05, **p < 0.01, ***p < 0.001, ****p < 0.0001.

**Supplementary Figure 9.**
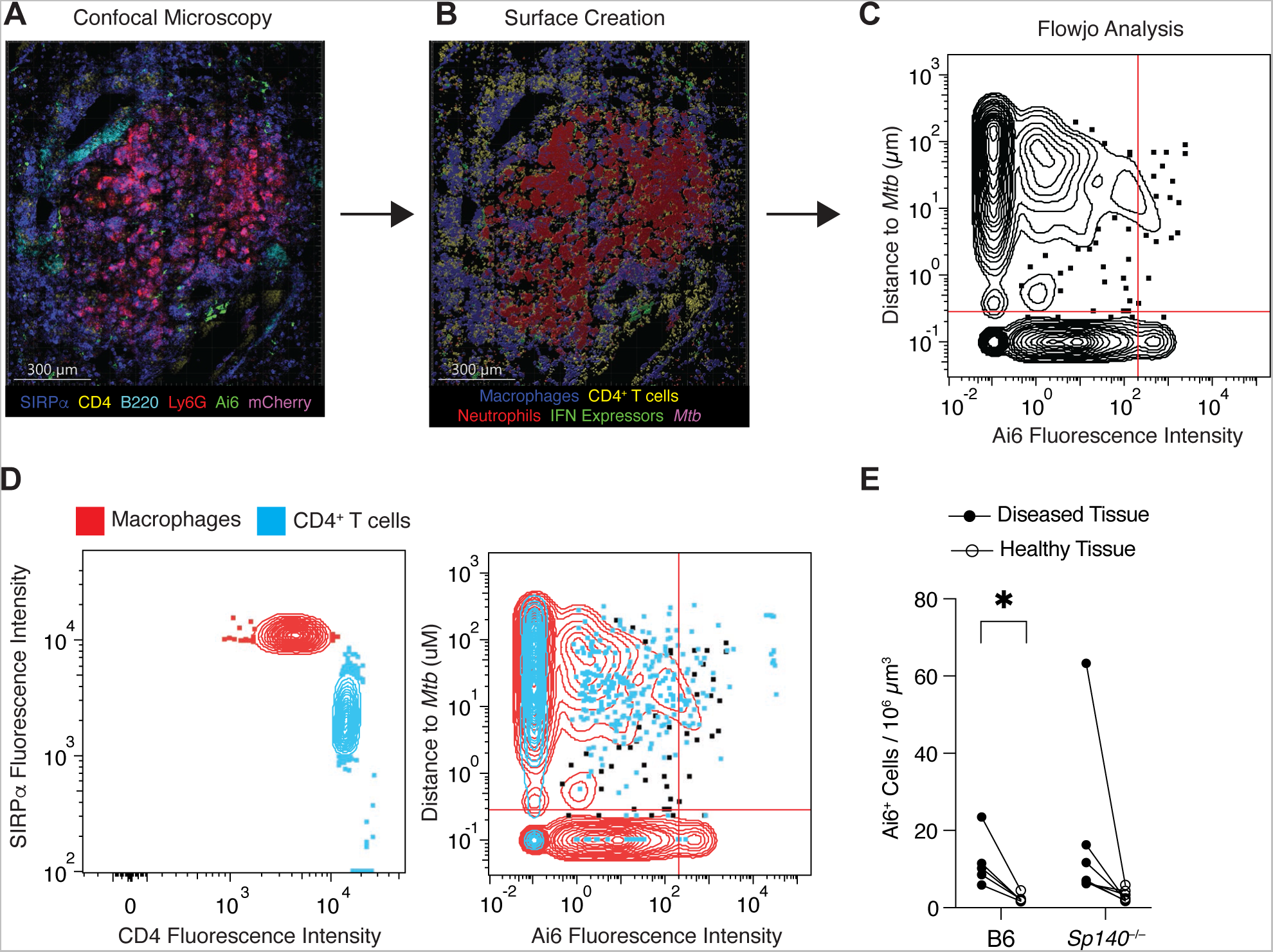
Histo-cytometry image analysis of *Mtb*-infected I-Tomcat Ai6 mouse lungs. (**A**) Representative confocal microscopy image of SIRPɑ (dark blue), CD4 (yellow), B220 (teal), Ly6G (red), Ai6 (green), and mCherry (magenta) staining in a section of mouse lung 25 days after *Mtb* infection. (**B**) Identifying cells within the image based on fluorescence intensity of individual channels as well as cellular morphology. (**C**) Flowjo analysis to quantify fluorescent signal intensity and distance to *Mtb* for cells identified in the image. (**D**) Representative Flowjo comparison of macrophages (red) and CD4^+^ T cells (blue) from an image of a diseased portion of an *Sp140*^−/−^ lung. (**E**) Number of Ai6^+^ cells per 10^6^ um^3^ in diseased (closed circle) and healthy tissue (open circle) from B6 (n = 5) and *Sp140*^−/−^ (n = 7) *Mtb*-infected mouse lungs. Mouse lungs were harvested 25 days after infection. Pooled data from two independent experiments are shown in (E). Statistical significance in (E) was calculated by multiple paired t tests. *p < 0.05.

**Supplementary Figure 10.**
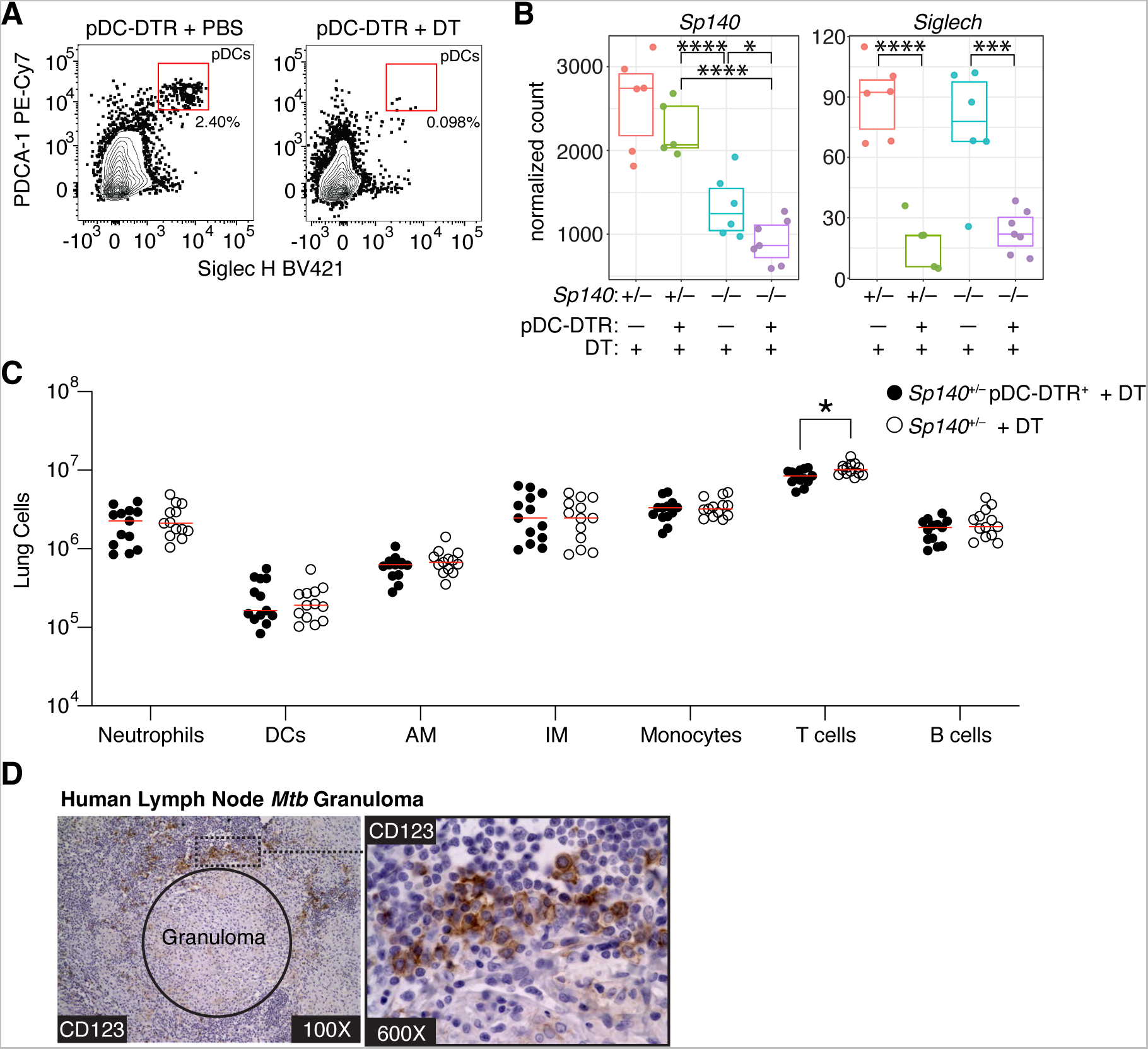
pDC-DTR mice specifically deplete pDCs without affecting major lung immune cell populations, and CD123 also identifies pDCs in *Mtb*-infected human lymph node samples. (**A**) Representative flow cytometry plot of splenic pDCs in *Sp140*^+/−^ pDC-DTR mice treated with PBS or DT from days 12 to 24 after *Mtb* infection. (**B**) RNA-sequencing validation of *Sp140* expression and pDC depletion efficiency based on *Siglech* expression in *Sp140*^+/−^ pDC-DTR mice treated with PBS or DT from days 12 to 24 after *Mtb* infection. (**C**) Number of various immune cell populations in *Mtb*-infected lungs of *Sp140*^+/−^ pDC-DTR (n = 13; filled circles) and *Sp140*^+/−^ mice (n = 13; open circles). Mice received DT from days 12 to 24 post-infection. (**D**) anti-CD123 (brown) and hematoxylin staining on *Mtb*-infected human lymph nodes. Mouse lungs and spleens were harvested 25 days after infection. Pooled data from two independent experiments are shown in (B) and (C). The bars in (C) represent the median. Statistical significance in (B) was calculated by the Wald test with multiple testing correction using the Benjamini and Hochberg method and in (C) by multiple unpaired t tests. *p < 0.05, ***p < 0.001, ****p < 0.0001.

**Supplementary Figure 11.**
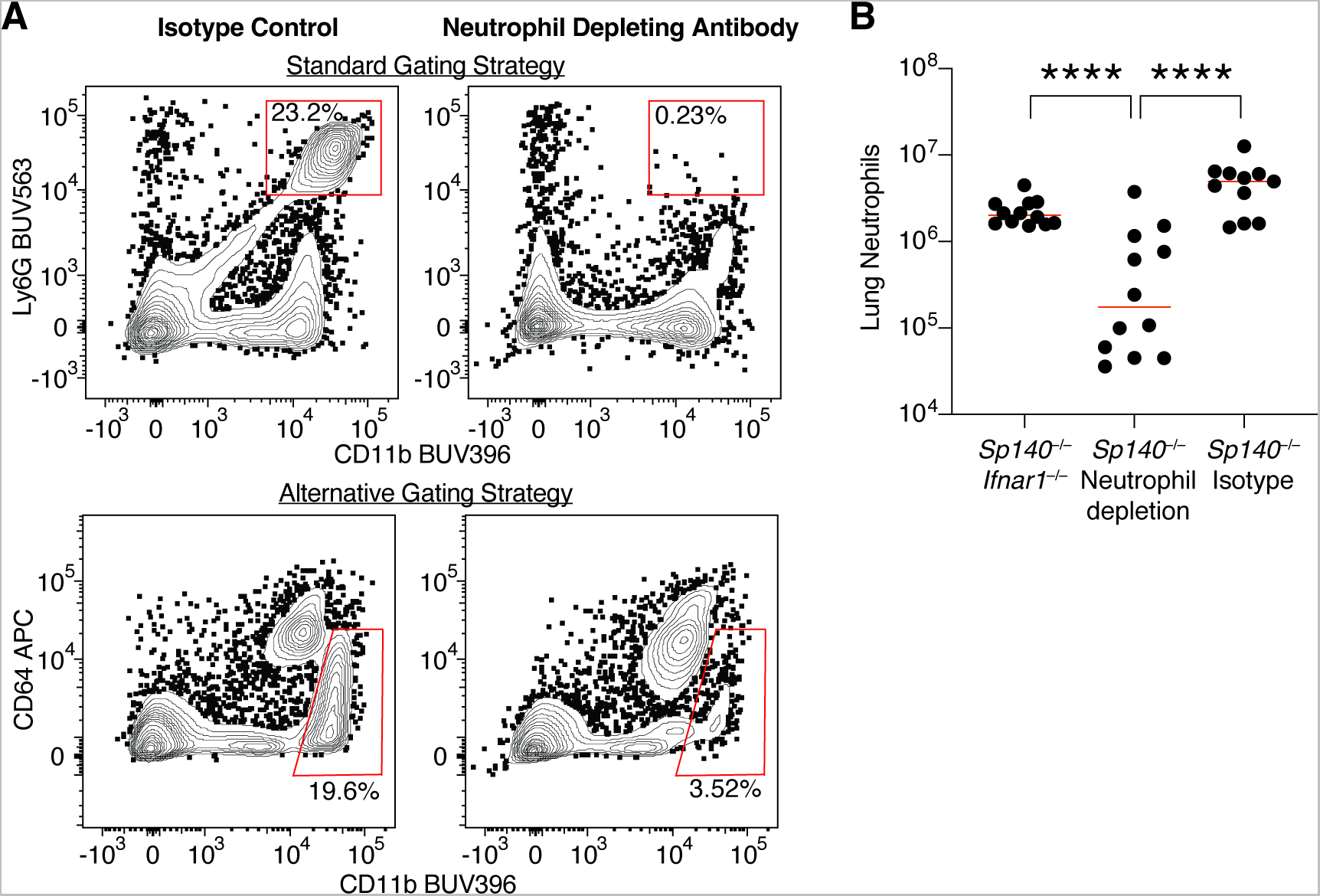
Anti-Ly6G antibody efficiently depletes neutrophils during *Mtb* infection. (**A**) Representative flow cytometry plots and (**B**) quantification of lung neutrophils in *Mtb*-infected *Sp140*^−/−^ mice treated with isotype control or clone 1A8 anti-Ly6G neutrophil depleting antibody from days 12 to 24 after *Mtb* infection. The standard neutrophil gating strategy uses anti-Ly6G clone 1A8, while the alternative strategy avoids anti-Ly6G antibody to confirm neutrophil depletion. Pooled data from two independent experiments are shown in (B). The bars in (B) represent the median. Statistical significance in (B) was calculated by one-way ANOVA with Tukey’s multiple comparison test. ****p < 0.0001

**Supplementary Figure 12.**
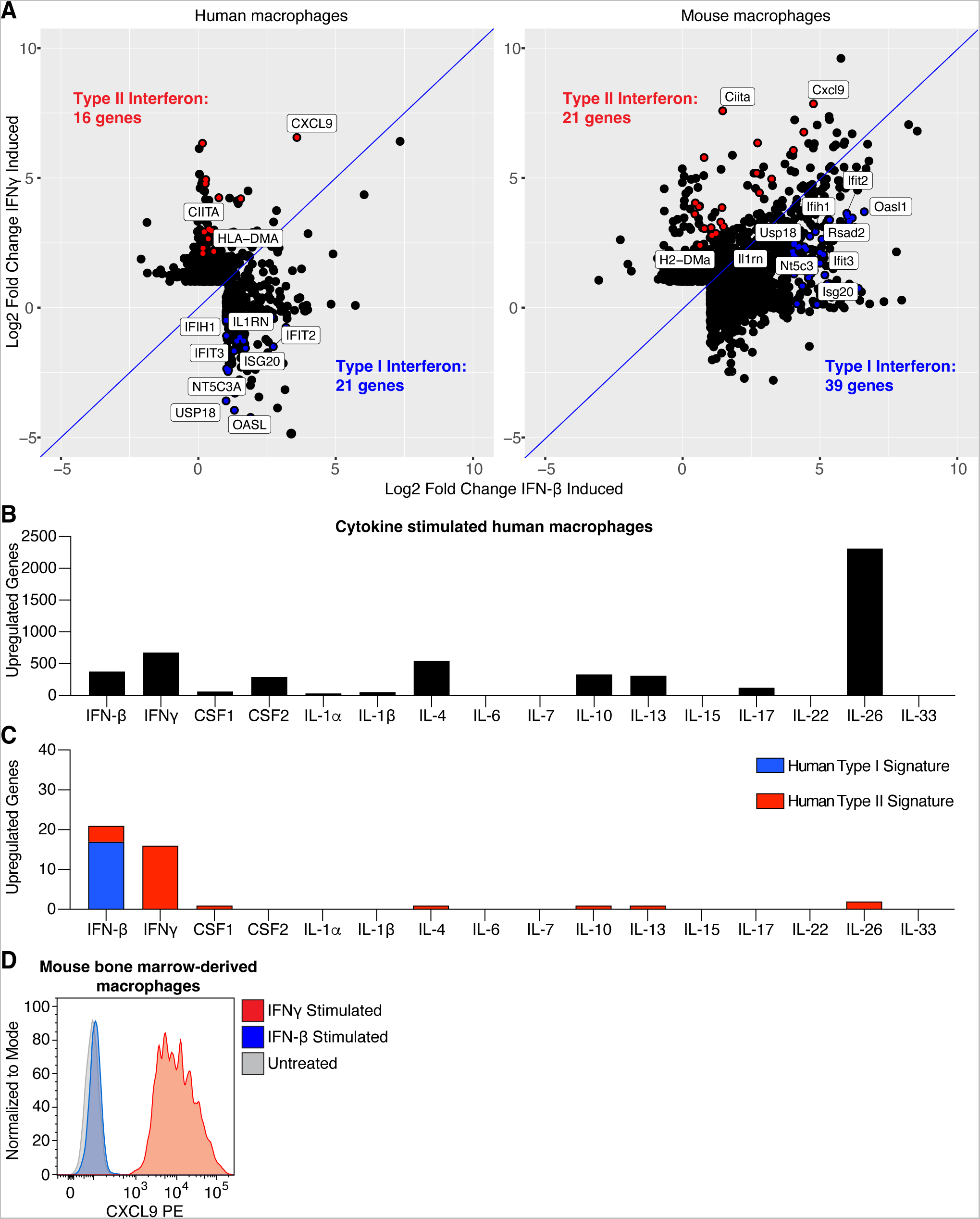
Generating gene signatures for identifying IFN*γ* and type I IFN responding cells. (**A**) Plots depicting the log_2_ fold change of genes upregulated in human macrophages or mouse bone-marrow derived macrophages following stimulation with IFN*γ* or IFN-β. Dots colored red indicate genes used for the IFN*γ* signaling gene signature while blue dots indicate those used for the gene signature for type I IFN responsiveness. Representative genes induced preferentially by IFN*γ* or IFN-β are labeled. (**B**) Number of genes upregulated in human macrophages following stimulation with each indicated cytokine and (**C**) Number of type I or II IFN gene signature genes induced after cytokine stimulation. (**D**) Representative flow cytometry histogram of CXCL9 expression by mouse bone marrow-derived macrophages that were untreated (grey), IFN-β stimulated (blue), or IFN*γ* stimulated (red).

**Supplementary Figure 13.**
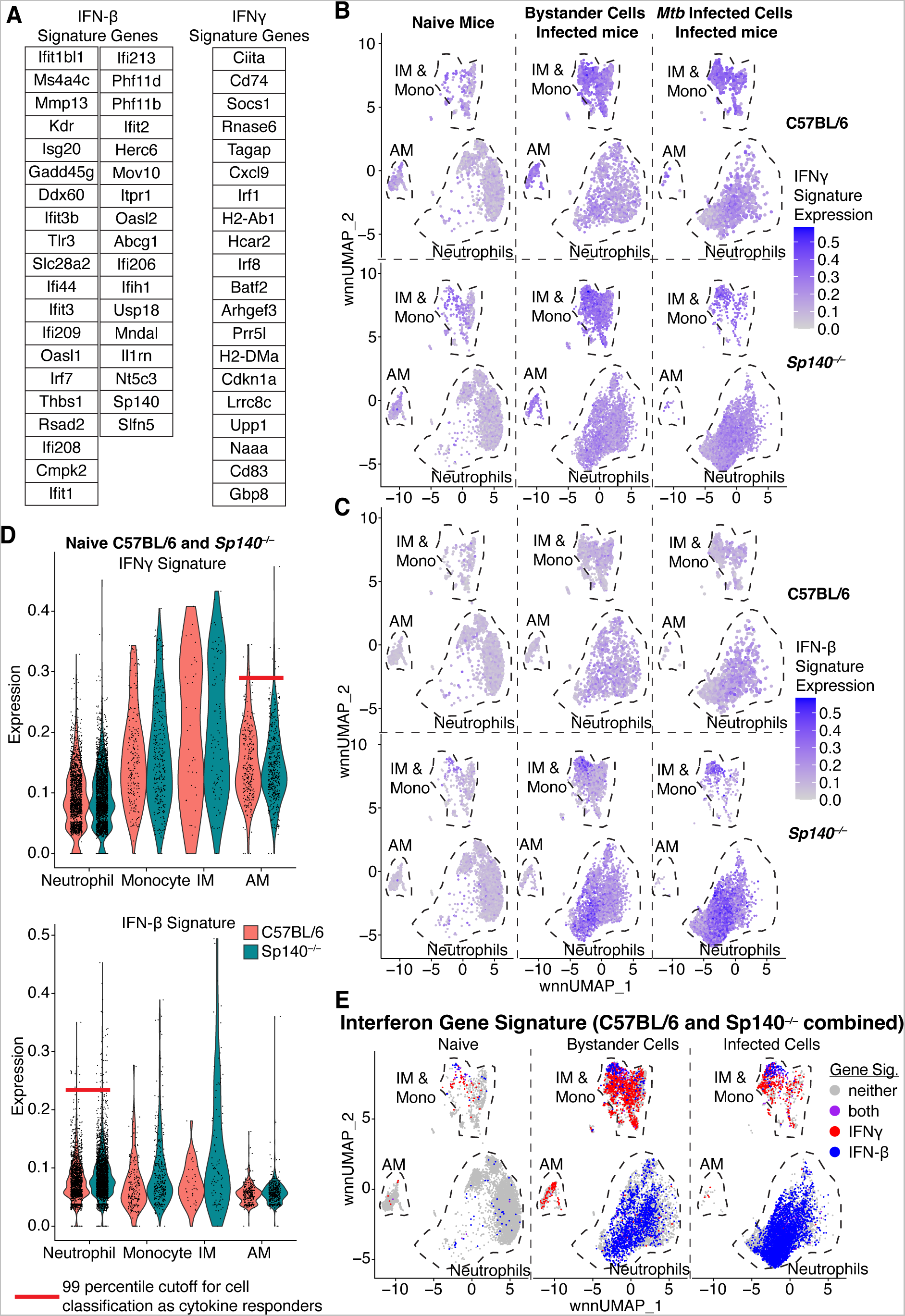
Applying gene signatures for identifying IFN*γ* and type I IFN responding cells to the myeloid scRNA-seq dataset. (**A**) List of genes from the cytokine stimulated mouse macrophages that were used for the gene signature for IFN*γ* or type I IFN responsiveness. (**B**) IFN*γ* gene signature or (**C**) type I IFN gene signature expression visualized by wnnUMAP plots of naïve, bystander, or *Mtb*-infected lung myeloid cells from B6 and *Sp140*^−/−^ mice. (**D**) Volcano plots of IFN*γ* and type I IFN gene signature expression on neutrophils, monocytes, IM, and AM from naïve B6 and *Sp140*^−/−^ mouse lungs. The red line indicates the 99% cutoff used to classify cells as type I or II IFN responders. (E) wnnUMAP plot comparing naïve, bystander, and *Mtb*-infected cells classified as cells that responded to type I IFN (blue), type II IFN (red), both (purple), or neither (grey) in lungs (B6 and *Sp140*^−/−^ combined).

**Supplementary Figure 14.**
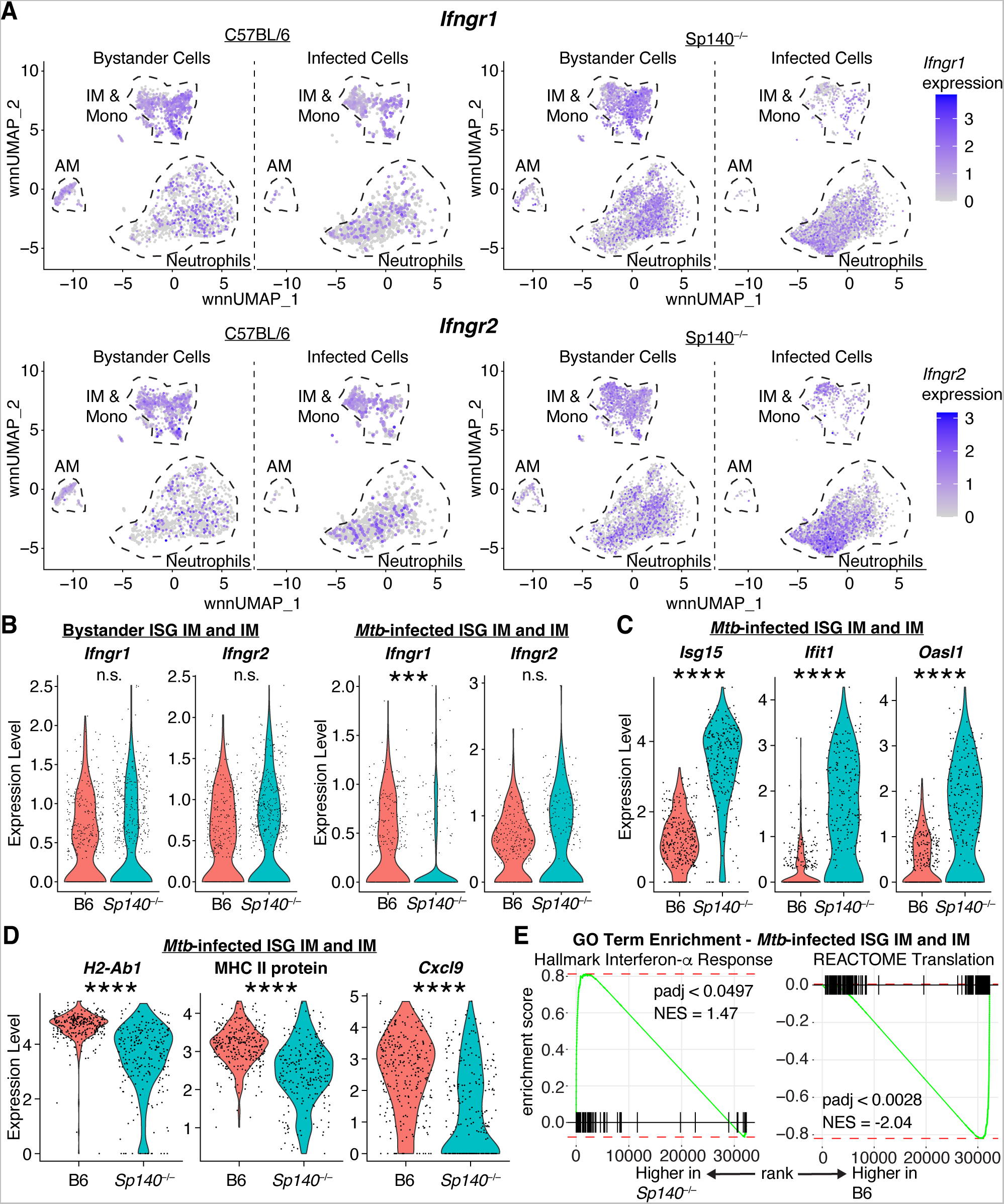
IFN*γ* receptor and IFN*γ* responsive genes are expressed at lower levels in *Mtb*-infected IMs in *Sp140*^−/−^ relative to B6 mice. (**A**) wnnUMAP plot of *Ifngr1* and *Ifngr2* expression on bystander and *Mtb*-infected cells from B6 and *Sp140*^−/−^ mice. (**B**) Volcano plots depicting expression of *Ifngr1* and *Ifngr2*, (**C**) representative type I IFN stimulated genes, including *Isg15*, *Ifit1*, and *Oasl1,* and (**D**) representative type II IFN stimulated genes, including *H2-Ab1* mRNA, MHC II protein, and *Cxcl9* mRNA on bystander and *Mtb*-infected IM and ISG^+^ IM from B6 and *Sp140*^−/−^ mice. (**E**) Gene ontology term enrichment of the Hallmark Interferon-a Response and the REACTOME Translation terms on *Mtb*-infected IM and ISG^+^ IM from B6 and *Sp140*^−/−^ mice. Statistical significance in (B), (C), and (D) was calculated by non-parametric Wilcoxon rank sum test with Bonferroni correction. ***p < 0.001, ****p < 0.0001.

**Supplementary Figure 15.**
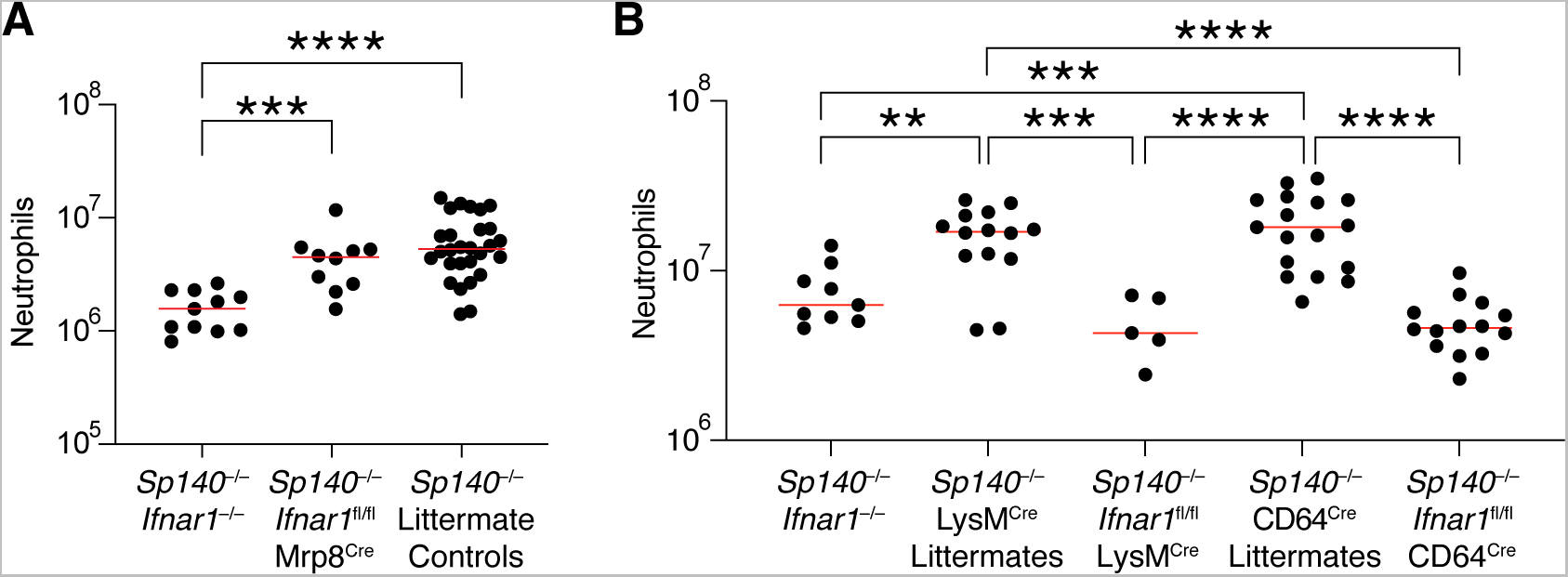
Lung neutrophil numbers are elevated in *Mtb*-infected *Sp140*^−/−^ mice with a specific deletion of type I IFN receptor on neutrophils but not on macrophages. (**A**) Lung neutrophil numbers in *Mtb*-infected *Sp140*^−/−^ *Ifnar1*^−/−^ (n = 11), *Sp140*^−/−^ *Ifnar1*^fl/fl^ Mrp8^cre^ (n = 10), and *Sp140*^−/−^ littermate control (n = 28) mice. (**B**) Lung neutrophil numbers in *Mtb*-infected *Sp140*^−/−^ *Ifnar1*^−/−^ (n = 9), *Sp140*^−/−^ LysM^cre^ littermate control (n = 14), *Sp140*^−/−^ *Ifnar1*^fl/fl^ LysM^cre^ (n = 5), *Sp140*^−/−^ CD64^cre^ littermate control (n = 20), and *Sp140*^−/−^ *Ifnar1*^fl/fl^ CD64^cre^ (n = 14) mice. The bars in (A) and (B) represent the median. Lungs were analyzed for the depicted experiments 24-26 days after *Mtb* infection. Pooled data from two independent experiments are shown. Statistical significance was calculated by one-way ANOVA with Tukey’s multiple comparison test. **p < 0.01, ***p < 0.001, ****p < 0.0001.

## Supplementary Tables

**Supplementary Table 1.**
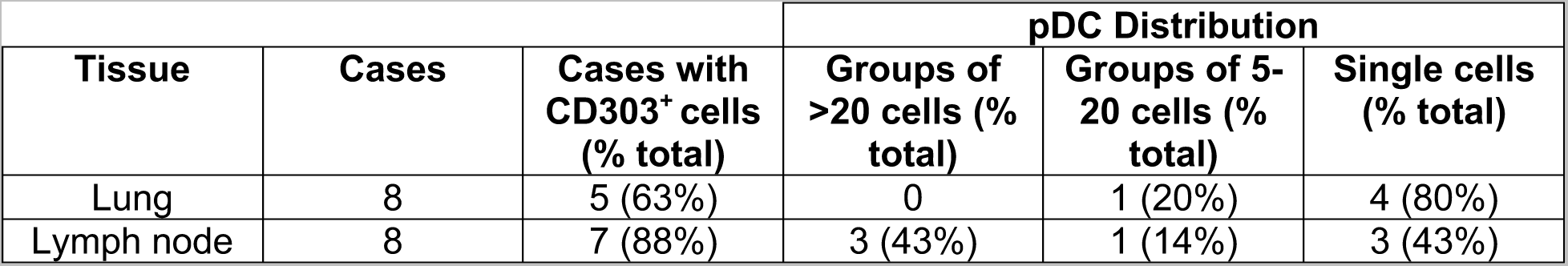
Quantification of the number of human lung and lymph nodes with pDCs in the same field of view as *Mtb* granulomas, as well as enumeration of the pDC distribution.

